# Aging and freezing of active nematic dynamics of cancer-associated fibroblasts by fibronectin matrix remodeling

**DOI:** 10.1101/2023.11.22.568216

**Authors:** Cécile Jacques, Louisiane Perrin, Joseph Ackermann, Samuel Bell, Olivier Zajac, Ambre Lapierre, Lucas Anger, Clément Hallopeau, Carlos Pérez-González, Lakshmi Balasubramaniam, Xavier Trepat, Benoît Ladoux, Ananyo Maitra, Raphael Voituriez, Danijela Matic Vignjevic

**Affiliations:** Institut Curie, PSL Research University, CNRS UMR 144; F-75005 Paris, France; Sorbonne Université and CNRS, Laboratoire Jean Perrin, F-75005, Paris, France; Laboratoire de Physique de l’Ecole normale supérieure, ENS, Université PSL, CNRS, Sorbonne Université, Université Paris Cité, F-75005 Paris, France; Université Paris Cité, CNRS, Institut Jacques Monod, F-75013 Paris, France; Institute for Bioengineering of Catalonia (IBEC), The Barcelona Institute for Science and Technology (BIST), 08028 Barcelona, Spain; Facutltat de Medicina, Universitat de Barcelona; 08036 Barcelona, Spain; Institució Catalana de Recerca i Estudis Avançats (ICREA); Barcelona, Spain; Centro de Investigación Biomédica en Red en Bioingeniería, Biomateriales y Nanomedicina (CIBER-BBN); 08028 Barcelona, Spain; Laboratoire de Physique Théorique et Modélisation, CNRS UMR 8089, CY Cergy Paris Université, F-95032 Cergy-Pontoise Cedex, France; INSERM, France

**Author notes:** Wellcome Trust / CRUK Gurdon Institute, University of Cambridge, Cambridge, UK. These authors contributed equally to this work (Cécile Jacques, Louisiane Perrin). These authors contributed equally to this work (Joseph Ackermann, Samuel Bell). Co-last and co-corresponding authors. Danijela Matic Vignjevic, Equipe Labellisée Ligue Contre le Cancer, Raphael Voituriez.

## Abstract

Cancer-associated fibroblasts (CAFs) are one of the most abundant cell types in tumor stroma. Exhibiting an elongated morphology, CAFs align with each other, closely resembling nematic ordering in liquid crystal physics. CAFs play a pivotal role in the genesis and remodeling of the extracellular matrix (ECM), with ECM proteins, especially fibronectin, reciprocally modulating CAF alignment and coherence. Yet, the intricate feedback loops between fibronectin deposition and CAF structuring remain largely unexplored. Here, we combined CAF live imaging, traction force microscopy, ECM microfabrication, and theoretical modeling to assess how the ECM influences the dynamics of nematically ordered CAFs. We found that CAFs dynamically orchestrate a fibronectin network that mirrors their nematic ordering. Over time, this passive nematic ordering of fibronectin, in turn, steers CAF rearrangement. Contrary to most cellular systems, where defects remain dynamic at a steady state, the ECM/CAF interplay profoundly alters the behavior of both CAF and ECM nematics, leading to aging-massive slowdown, and even freezing of defect dynamics. This results in a CAFs capsule where aligned areas and defects are spatially and temporally fixed, yet active, exerting forces at the substate and transmitting forces between cells. Functionally, while defects may be permissive, it is the fibronectin loss–induced fluidization of the CAF capsule that critically undermines its barrier function.

## Introduction

The most abundant cell type in the tumor stroma is cancer-associated fibroblasts (CAFs) ^1, 2^. In many tumor types, particularly in colorectal, breast, and pancreatic cancers, CAFs and the extracellular matrix (ECM) form a protective capsule that envelops tumors ^3^, and plays a key role in regulating their evolution. Contrary to the belief that this capsule acts passively to prevent tumor growth ^4–6^, we have recently found that CAFs exert active compression on cancer cells, thereby triggering mechanical signaling that slows down their proliferation ^3^.

The ECM is mainly produced and remodeled by CAFs ^7, 8^. In return, ECM proteins, in particular fibronectin, facilitate communication among fibroblasts ^9–11^. This intercellular communication favors alignment within the fibroblasts, and coordinates CAFs’ supracellular contractility and dynamics ^3^. This results in a reciprocal relationship between ECM-induced cell activities and cell-driven ECM alterations ^10, 12^, which were also reported for other cell types at the single cell level ^13–15^. Yet, the impact of this interaction on the integrated organization of the ECM/CAF capsule and tumor progression is still to be understood.

CAFs have an elongated shape and tend to align with each other in a process termed “cell collision guidance” ^16^. In liquid crystal physics, the alignment of the long axis of generic apolar elongated particles is described as nematic ordering ^17–19^. Nematic order inherently involves “topological defects” (induced by geometric frustration, thermal fluctuations or activity), which are singular points in space where particles with different orientations meet leading to an ill-defined local alignment ^17, 19^. Recent studies have demonstrated that substrate anisotropy or topographical patterns—such as ridges—can impose nematic order and stabilize defects in cell monolayers ^20–23^. However, these approaches rely on external cues to direct alignment and arrest defect motion. In cellular monolayers, defects have been described as spots where biologically important cellular behaviors such as extrusion, multilayering and/or polarity changes are favored ^24–27^. Therefore, while the aligned CAFs within the capsule create a barrier that prevents tumor growth and spread, defects in the alignment could provide weak points, allowing cancer cells to escape. In this context, how the ECM/CAF interaction controls the nematic organization and defect dynamics of CAF monolayers and what consequences these dynamics can have on biological functions such as cancer cell dissemination remains unknown.

Here, we uncover a distinct mechanism of defect arrest that emerges intrinsically from ECM/CAF feedback and monolayer aging, rather than imposed substrate cues. In this study, we combined CAFs live imaging over a long period with traction and stress force microscopy, ECM patterning, and theoretical modeling to assess how the ECM influences the dynamics of CAFs’ nematic alignment. We found that CAFs dynamically orchestrate a fibronectin network that mirrors their nematic ordering. In response and over time, the passive nematic ordering of fibronectin guides the organization of CAFs. While prior work has shown that collagen remodeling can align fibroblasts ^28^, our results reveal that ECM/CAF feedback drives a unique aging process, leading to dynamic arrest of defects without external guidance. Note that here, by aging, we mean that the process is non-stationary, but slowly varying in time. In contrast to most cellular systems described so far, and more generally of active nematic systems, in which defects have a finite density and remain dynamic even in steady state - they move, emerge, and annihilate over time ^17–19, 25^ - we show that the ECM/CAF feedback interaction dramatically impacts the dynamics of both the CAFs and ECM nematic fields, leading to massive slow down and even freezing of defect dynamics. As a result, both the aligned areas and topological defects within CAFs and ECM became stationary in both space and time but remained active, with in particular larger traction forces focused at defects, potentially weakening the CAF layer and favoring cancer cell dissemination.

## Results

### CAFs and the fibronectin matrix they produce establish layers with congruent nematic ordering

To investigate whether CAFs exhibit nematic ordering similar to other elongated cells, such as myoblasts ^29–31^, we seeded CAFs isolated from human colorectal tumors on 11 kPa polyacrylamide (PAA) gels coated with monomeric collagen I to maintain physiological substrate rigidity ^32^. Over time, CAFs established a monolayer, comprising large regions of aligned cells that formed a nematic order (**Fig. 1A, Supp. Fig.1 A, B**). This nematic ordering was disrupted locally by topological defects, with expected charges of +1/2 and -1/2, and characteristic shapes resembling comets or triangles, respectively. Both defect charges were observed with a similar density of 1.5 defects/mm² within the CAF layer (**Fig. 1B, Supp. Fig. 1C**).

**Figure 1.**
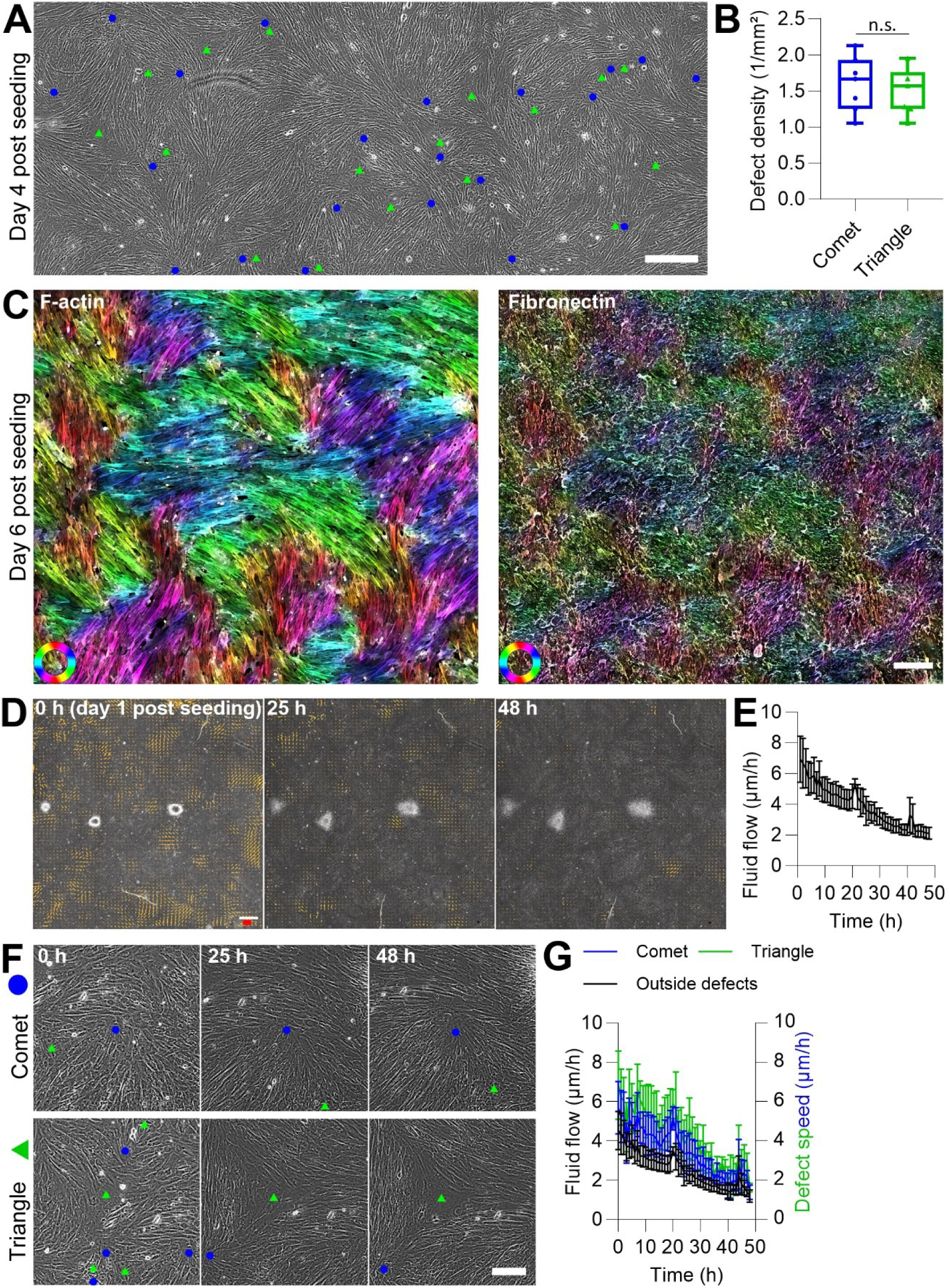
CAFs and fibronectin form highly correlated nematic ordered layers with stationary defects in the CAF layer. A) Bright field of a CAF layer cultured on an 11kPa PAA gel for 4 days. Automated detection of topological defects: +1/2 defects or comet defects (blue dots) and -1/2 defects or triangle defects (green triangles). Scale bar: 400 µm. B) Defect density in the nematically ordered CAF layer. Comet and triangle defects are represented in blue and green, respectively. 7 fields of view from 3 independent experiments, unpaired two-sided t-test: p-value = 0.632. Boxplot: middle bar= median, edges bars= 25th and 75th percentiles, whiskers= extent of data. C) Left: Orientation of CAFs based on F-actin staining (phalloidin); right: orientation of fibronectin network, 6 days after seeding. Cells or fibers with the same orientation are represented with the same color. Colored circle: orientation colormap. Scale bar: 200 µm. D) Bottom: Bright field time-lapse imaging of the CAF layer and evolution of the velocity field over time. Time-lapse started one day after seeding. Orange arrows represent the local velocity vectors. Red scale bar: 400 µm/h. White scale bar: 500 µm. E) Evolution of the mean of the velocity field of the CAF layer over time. The black line represents the mean of 3 independent experiments, and the error bars represent the standard error of the mean. F) Representative automated detection of comet (top, blue circle) and triangle (bottom, green triangle) defects over time. Scale bar: 100 µm. G) Defect speed over time in the nematically ordered CAF layer. Comet and triangle defects are represented in blue and green, respectively. Areas outside defects are represented in black. The lines represent the mean of 3 independent experiments, and the error bars represent the standard error of the mean.

To examine how these defects influence the deposition and organization of fibronectin by CAFs, we assessed a correlation between the orientation of CAFs (based on F-actin staining) and fibronectin once cell monolayer was confluent and nematic order fully established, approximately 6 days after cell seeding (**Fig. 1C**). The comparative analysis of orientation maps revealed a similar local orientation between CAFs and fibronectin layers, with a 92± 5% local correlation of fibers’ orientation (4 independent experiments). Even though cell density and fibronectin deposition were heterogeneous, we did not observe preferential accumulation of either at defects or in aligned areas (**Supp. Fig. 2A, B**). Of note, CAFs deposited a very small amount of collagen I (**Supp. Fig. 2C**), as previously shown ^3^, indicating that fibronectin is the predominant ECM component deposited during the course of the experiment.

Altogether, this data shows that the deposited fibronectin fibers almost perfectly reproduce the nematic ordering of the CAFs layer.

### Freezing of the CAFs active nematic is achieved after transient, aging dynamics

Typically, active nematics in both biological materials, such as cell monolayers and artificial or in silico systems, can reach a chaotic stationary state in which defects emerge, move and annihilate randomly ^17–19^. If the orientation of fibronectin deposition matches the local CAFs orientation consistently across space and time, then over several days, with CAFs changing dynamically, the cumulative effect of these random orientation signals should produce a fibronectin layer with uniform density and randomly oriented fibers. Because our observations contradicted this prediction, we questioned whether the anticipated chaotic stationary dynamics was occurring within the CAFs’ layer. To investigate the dynamics of the defects, we carried out time-lapse imaging of the CAFs’ layer starting 2 days after cell seeding. Over a period of 15 hours, we observed no changes in either the position or shape of the defects (**Supp. Fig. 3A**). By tracking of the defects’ core positions, we generated trajectories relative to the size of the defect, which provides an estimate of the error in determining the position of the defects. We found that the defect’s total displacement was always significantly smaller than the typical defect’s size, and thus within measurement noise (**Supp. Fig. 3B**); indicating no significant motion of defects beyond fluctuations. Similar nematic alignment (**Supp. Fig. 4A**) and arrested defect dynamics (**Supp. Fig. 4B**) were also observed for fibroblasts isolated from normal tissue adjacent to tumor (normal-associated fibroblasts, NAFs) suggesting that these behaviors are not restricted to a cancer context.

This arrested dynamic is in striking contrast with the dynamic defects observed in various active nematic systems, including cell monolayers ^33^. A natural question arises regarding the timeline of this process. To analyze the transient dynamics leading to the freezing of the CAF monolayer, we performed time-lapse imaging of the entire CAF layer starting one day after seeding (**Fig. 1D**). The quantification of the evolution of cellular flows by PIV over 48 hours showed a global slowdown - and almost complete arrest of cellular flows after 3 days (**Fig. 1E**). A comparable kinetic slowdown was also observed for the speed of both types of defects, as well as for cellular flows in areas empty of defects (**Fig. 1F, G**), indicating a spatially uniform, scale-independent aging process. This observed freezing of the flow over time was not dependent on substrate stiffness (**Supp. Fig. 5A**) or a specific ECM coating (**Supp. Fig. 5B**).

Taken together, these results show that the dynamics of defects within the CAFs layer exhibit slowdown, or aging, which eventually leads to an arrested state.

### In the nematic-ordered layer, a majority of CAFs remain stationary yet actively exert and transmit forces

We next wondered whether the stasis we observed was limited to defects or if it also applies to the individual CAFs within the nematic ordered layer. In other words, was the whole system frozen? To monitor the movement of individual CAFs, we genetically modified the CAFs to express a fluorescent marker (LifeAct-GFP), mixed 20% of these CAFs with unlabeled CAFs and started time-lapse imaging after nematic order was established. Tracking of the labeled CAFs over a 15-hour period revealed two distinct types of movements within the monolayer (**Fig. 2A**). While a few CAFs showed directed, linear movement with higher velocities of around 6 µm/h and persistence above 1 (see M&M for definition), the majority of CAFs (92% of n=199) appeared to be fluctuating around their initial positions, with no preferred direction of motion. These fluctuating cells displayed slower typical speeds of approximately 1 µm/h, and a persistence below 1, consistent with random motion (**Fig. 2B; Supp. Fig. 6A**). Similar cell dynamics were also observed for NAFs (**Supp. Fig. 4C, D**). This data shows global freezing of the cellular flow within the full monolayer and not only of defects.

**Figure 2.**
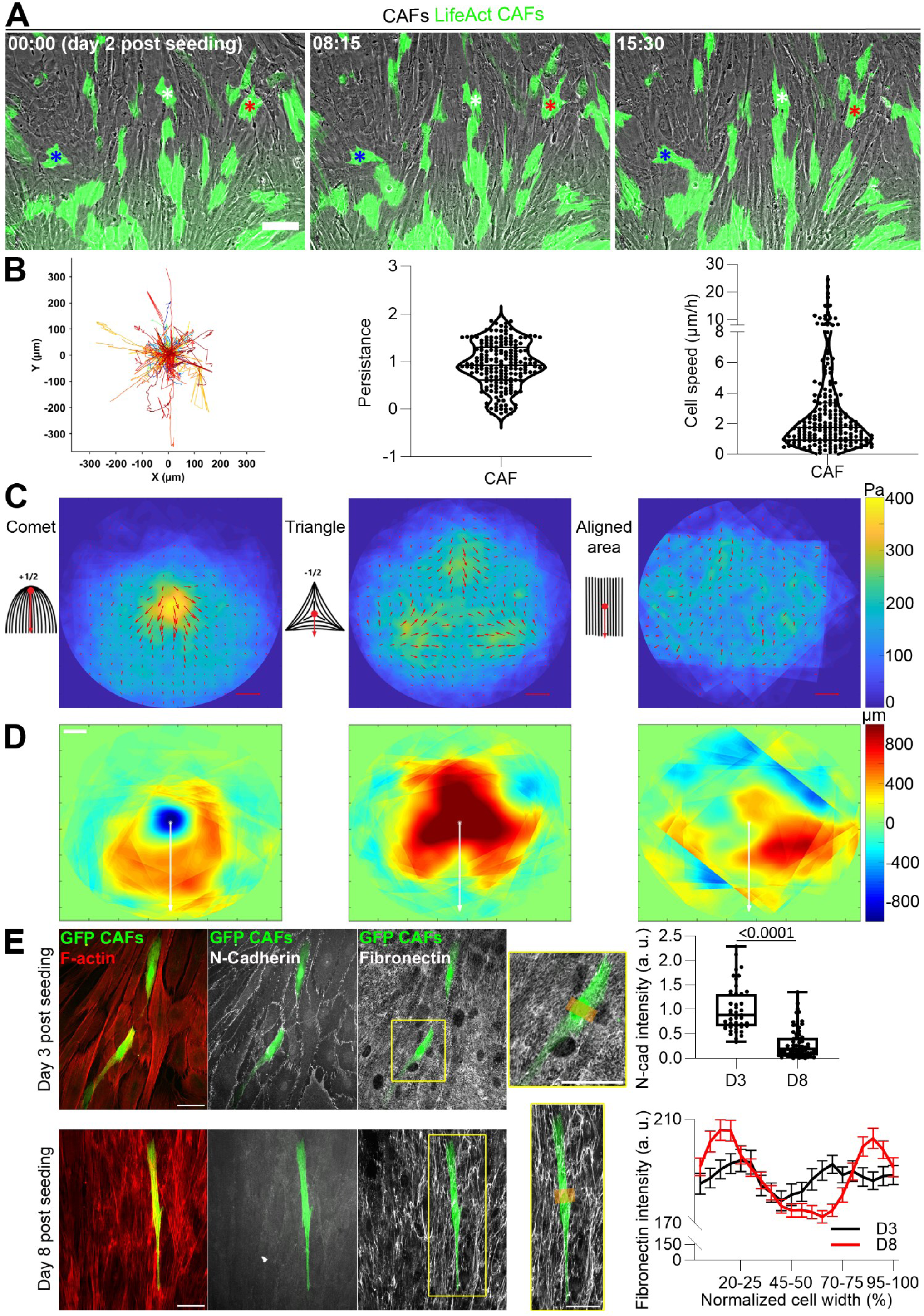
CAFs in the nematic ordered layer are immobile. A) Time-lapse imaging of LifeAct-GFP expressing CAFs (green) mixed with unlabeled CAFs 2 days after seeding. Blue, white, and red stars follow the position of three different CAFs over time. Scale bar: 100 µm. B) Left: CAFs trajectories in the whole field of view. Each trajectory represents one CAF, 199 trajectories from 2 independent experiments. See also supplementary figure 6A. Right: Quantification of CAFs persistence and speed. Each dot represents one CAF, 199 cells from 2 independent experiments. C) Schemes representing the core and the direction used to average aligned areas, comet, and triangle defects. Average of traction force magnitude maps (colormap) and vectors (red arrows) for aligned areas, comet, and triangle defects. Red scale bar: 800 Pa. 50 comet defects, 42 triangle defects, and 18 aligned areas from 4 independent experiments. D) Average of isotropic stress maps for aligned areas, comet, and triangle defects. White stars and vectors correspond to the red dot and vectors of the respective scheme in (C). White scale bar: 100 µm. 50 comet defects, 42 triangle defects, and 18 aligned areas from 4 independent experiments. E) Left: Layer of unlabeled CAFs mixed with GFP CAFs (green) fixed at 3 or 8 days post seeding, and stained for F-actin (phalloidin, red), N-cadherin (gray), and fibronectin (gray). Insets show the regions (orange boxes) used for quantification of the fibronectin intensity. Scale bars: 50 µm. Right: For N-cadherin, each dot represents one cell. 45 and 65 cells were analyzed at day 3 and 8, respectively, from 3 independent experiments; Mann Whitney test. Boxplot: middle bar= median, edges bars= 25th and 75th percentiles, whiskers= extent of data. For fibronectin, the quantification was performed on 51 and 134 cells at day 3 (black line) and 8 (red line), respectively. The lines represent the mean of 3 independent experiments, and the error bars represent the standard error of the mean.

Next, we wondered if only cell motility was decreased or if CAFs, in general, became less active. As CAFs are highly contractile cells ^8, 10, 34^, we examined if they remain contractile even in stalled defects. To assess both the force patterns generated by the CAF layer on the substrate and the internal stress in the monolayer, we performed traction force and internal stress microscopy (**Fig. 2C, D**). For each defect topology, multiple samples were collectively averaged, using their core and direction to align them consistently (for triangle defects, one of the three equivalent directions was used). While low levels of traction forces and internal isotropic stress without a discernible pattern were observed in aligned areas, defects show specific force patterns and isotropic stress patterns similar to other contractile cellular systems that are not stalled ^17, 26^. In the case of comet defects, traction forces were highest at the head of the comet core, with force directions pointing towards the tail of the comet, with the head of the comet consistently showing compression (negative isotropic stress) and the tail displaying tension (positive isotropic stress). For triangle defects, peaks of traction forces were evident at each triangle vertex with forces directed towards the triangle core; with the center of the triangle under tension (positive isotropic stress). Thus, even though CAFs and defects in their nematic layer are frozen, they are still active as cells continue to exert forces, behaving globally as an active contractile monolayer.

Such distinctive patterns at defects could potentially stem from the supracellular units formed between CAFs. Indeed, the low traction forces observed in the aligned area suggest that forces are transmitted between the CAFs. As the comet core and triangle vertices are the termini of aligned areas, the transmission of forces between CAFs is abolished, and forces are transmitted to the substrate, resulting in the visible peaks of traction in both cases.

A possible candidate for such interconnections between CAFs is N-cadherin, a key cell-cell adhesion protein in mesenchymal cells ^35^. Alternatively, CAFs could be linked via stitch adhesions with fibronectin acting as a glue ^9, 11^. To examine how CAFs were connected, we stained for N-cadherin and fibronectin in CAF layers of varying densities (**Fig. 2E**). At lower densities, where CAFs were more spread out and less aligned, we observed N-cadherin at cell-cell junctions. However, as cell density increased, leading to elongated, nematic-ordered CAFs, N-cadherin presence diminished. Conversely, fibronectin is enriched at the cell periphery as the CAF density increased. These findings suggest that in a nematic ordered layer, CAFs may be indirectly interacting via fibronectin.

Therefore, the observed lack of directed movement of both defects and individual CAFs within nematic layers could be attributable to fibronectin depositions interposed between CAFs and the substrate, potentially immobilizing the system.

### Fibronectin depletion restores the dynamics of cells and defects in CAF nematic layer

To assess whether the immobilization of the CAFs layer dynamics was indeed a result of fibronectin deposition, we depleted fibronectin in CAFs using siRNA (**Fig. 3A**) and performed time-lapse imaging (**Fig. 3B**). By tracking individual cells using Hoechst labeling of nuclei, we observed that fibronectin-depleted cells (siFN) exhibited longer and straighter trajectories (**Fig. 3C**), and moved with greater persistence and speeds (**Fig. 3D; Supp. Fig. 6B**), compared to CAFs treated with control siRNA (siCtrl). Thus, upon depletion of fibronectin, the mobility of CAFs within the layer significantly increased; this points towards a direct role of FN deposition in the observed aging dynamics, and excludes a density-dependent jamming phenomenon only (discussed in ^36^).

**Figure 3.**
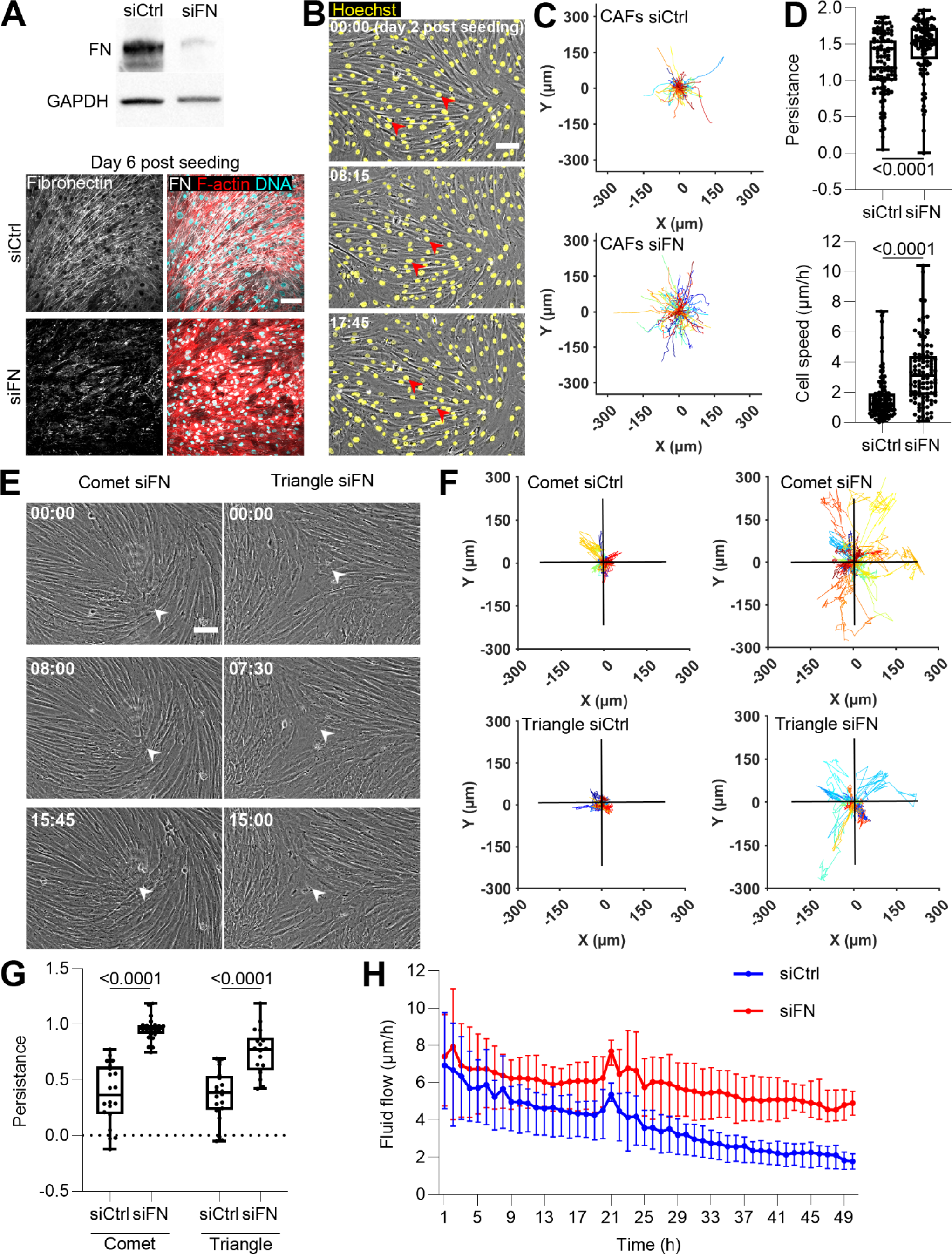
Dynamics of defects and CAFs is restored upon fibronectin depletion. A) Depletion of fibronectin in CAFs evaluated by western blot (top) and immunofluorescence (bottom). Cells were transfected with either control siRNA (siCtrl) or siRNA against fibronectin (siFN). GAPDH was used as a loading control. Cells were stained for F-actin (phalloidin, red), DNA (DAPI, cyan), and fibronectin (white) 6 days post seeding. Scale bar: 100 µm. B) Bright field time-lapse imaging of CAFs transfected with siRNA against fibronectin (siFN) and labeled with Hoechst (yellow) 2 days after seeding. Arrowheads follow CAFs over time. Time, hours:minutes. Scale bar: 100 µm. C) Trajectories of CAFs transfected with either control siRNA (siCtrl) or siRNA against fibronectin (siFN). Each trajectory represents one CAF; 100 CAFs for each condition from one experiment. See also supplementary figure 6B. D) Quantification of persistence (top) and speed (bottom) of CAFs transfected with either control siRNA (siCtrl) or siRNA against fibronectin (siFN). Each dot represents one CAF; 100 CAFs for each condition from one experiment; two-tailed unpaired t-test: p-values mentioned on the graph. Boxplot: middle bar= median, edges bars= 25th and 75th percentiles, whiskers= extent of data. E) Bright field time-lapse imaging of defect dynamics in fibronectin-depleted CAF layers (siFN) 2 days after seeding. White arrows follow the defect core over time. Time, hours:minutes. Scale bar: 100 µm. F) Trajectories of defect cores in control (siCtrl) and fibronectin-depleted (siFN) CAF layers. Each trajectory represents one defect. siCtrl: 20 comet and 20 triangle defects; siFN: 24 comet and 18 triangle defects from at least 3 independent experiments. Horizontal and vertical black lines represent the defect size (400 µm). See also supplementary figure 6C. G) Quantification of the persistence of defects in control (siCtrl) and fibronectin-depleted (siFN) CAFs layers. Each dot represents one defect; siCtrl: 20 comet and 20 triangle defects; siFN: 24 comet and 18 triangle defects from 5 independent experiments; two-tailed unpaired t-test: p-values mentioned in the graph. Boxplot: middle bar= median, edges bars= 25th and 75th percentiles, whiskers= extent of data. H) Evolution of the mean of the velocity field of control (siCtrl, blue) and fibronectin-depleted (siFN, red) CAF layer over time. The time-lapse started one day after seeding. The line represents the mean of 3 independent experiments, and the error bars represent the standard error of the mean.

Subsequently, we assessed whether the increased motility of CAFs in the absence of fibronectin also influenced the dynamics of defects in the CAFs’ nematic layer. To that end, we performed time-lapse imaging of CAFs treated either with siCtrl or siFN (**Fig. 3E**). Tracking of the cores of defects revealed that in the absence of fibronectin, defects become persistent (**Fig. 3F, G; Supp. Fig. 6C**). Indeed, while defects in the layer of control CAFs fluctuated around their initial position (**Fig. 3F**), defects in the fibronectin-depleted CAFs layer exhibited longer and straighter trajectories (**Fig. 3F**), moving with greater persistence (**Fig. 3G; Supp. Fig. 6C**). In particular, comet defects were motile, moving along their head-tail direction, indicating that fibronectin-depleted CAFs were forming a contractile nematic system ^18, 26^. Triangle defects, in contrast, displayed lower speeds and persistence compared to comet defects, likely due to a lack of a preferred migration direction^24^, as expected in active nematics.

Finally, for fibronectin-depleted CAFs, the slowdown of cellular flows was significantly less pronounced and did not lead to freezing observed in control CAFs (**Fig. 3H**).

Therefore, the fact that both individual cells and defects remain more dynamic in the absence of fibronectin suggests that fibronectin deposition is causing the freezing or aging, of the system.

### Established fibronectin patterns freeze CAF nematic dynamics and migration of individual cells

Until now, we showed that CAF nematic ordering generated fibronectin patterns with the same nematic order and that fibronectin deposition was necessary for freezing the system. To further characterize the mechanism of freezing, we tested if the interaction of CAFs with a pre-existing fibronectin pattern could be sufficient to induce the freezing of the dynamics of the CAF layer. For this, we cultured CAFs on PAA gels, allowing them to create nematic ordering (**Fig. 4A, B**; CAFs 1). Subsequently, we removed the CAFs to obtain the corresponding fibronectin patterns with nematic ordering (**Fig. 4A, B**; FN 1). On those fibronectin patterns, we then seeded new CAFs. Once these CAFs formed a confluent layer, we determined their organization as well as the organization of the newly deposited fibronectin (**Fig. 4A, B**; CAFs 2 and FN 2). The comparative analysis of orientation maps revealed a similar local orientation between both fibronectin patterns and both CAFs layers (**Fig. 4C**). These data show that the fibronectin pattern was instrumental in instructing the CAFs to align into its specific nematic ordering configuration.

**Figure 4.**
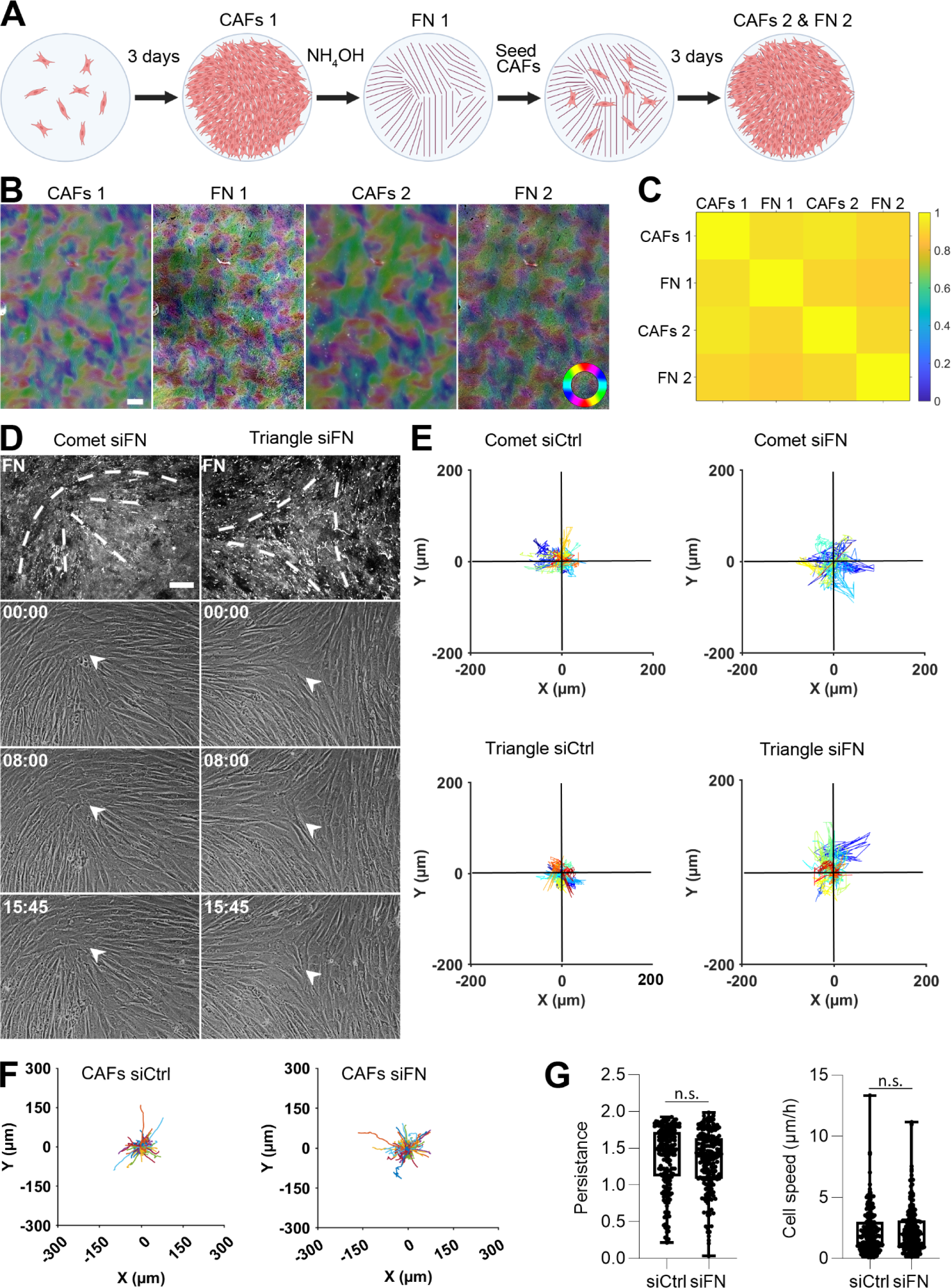
Fibronectin patterns freeze CAF nematics. A) Scheme representing the experiment design: CAFs were seeded on 11kPa PAA gel and cultured for three days. CAFs were imaged (CAFs 1) and then removed using NH_4_OH, before staining and imaging of the deposited fibronectin (FN 1). On those fibronectin patterns, new CAFs were seeded and cultured for an additional three days. CAFs were imaged (CAFs 2) and then removed using NH_4_OH, before staining and imaging of the deposited fibronectin (FN 2, representing the sum of FN1 and newly deposited fibronectin). B) Orientation map of CAFs (CAFs 1 and 2) and fibronectin layers (FN 1 and 2). Colored circle: orientation colormap. Scale bar: 600 µm. C) Mean of the local spatial orientation correlation between the CAFs and fibronectin layers. Results from 3 independent experiments. D) Fibronectin (FN, gray) stained with specific antibodies, highlights defects - comet (left) and a triangle (right). White dashed lines represent the shape of a comet (left) and a triangle (right). Bright-field time-lapse imaging of defect dynamics in fibronectin-depleted CAFs layers (siFN) seeded on preformed fibronectin patterns 2 days before imaging. White arrowheads follow the positions of defects’ cores over time. Time, hours:minutes. Scale bar: 100 µm. E) Trajectories of defect cores in control (siCtrl) and fibronectin-depleted (siFN) CAFs layers seeded on preformed fibronectin patterns. Horizontal and vertical black lines represent defect size (400 μm). Each trajectory represents one defect; siCtrl: 18 comets and 12 triangles, siFN: 17 comets and 15 triangles from 3 independent experiments. F) Trajectories of control CAFs (siCtrl) and fibronectin-depleted CAFs (siFN) seeded on preformed fibronectin patterns 2 days before imaging. Each trajectory represents one CAF (200 cells for each condition, from 3 independent experiments). G) Quantification of persistence and velocity of CAFs transfected with either control siRNA (siCtrl) or siRNA against fibronectin (siFN) and seeded on a fibronectin pattern. Each dot represents one CAF (200 cells for each condition, from 3 independent experiments); two-sided unpaired t-test p-value = 0.3527 and 0.4443. Boxplot: middle bar= median, edges bars= 25th and 75th percentiles, whiskers= extent of data, red cross= outliers.

Next, we explored if fibronectin patterns or, alternatively, fibronectin depositions between CAFs, were responsible for freezing CAF nematics. To discriminate between these possibilities, we produced fibronectin nematic patterns as described above and seeded either control CAFs (siCtrl) or fibronectin-depleted CAFs (siFN) on top of them. As observed above, CAFs aligned according to the fibronectin nematics and defects (**Fig. 4D**). Over a 15-hour period, no differences were observed in defect position (**Fig. 4E**). Indeed, both, control and fibronectin-depleted CAFs fluctuated around their position, with a fluctuation range smaller than the defects’ size (**Fig. 4E**). Accordingly, cellular flows were reduced on anisotropic fibronectin patterns compared to collagen I; as well as on isotropic fibronectin fibers (**Supp. Fig. 7**). By tracking individual CAFs within the nematic layer, we found that fibronectin patterns were also sufficient to arrest the migration of individual cells, independently of whether they produce fibronectin (**Fig. 4F, G**). While a subset of CAFs followed the orientation of fibronectin patterns, direction of migration was not affected by the local fibronectin density (**Supp. Fig. 8**).

Therefore, fibronectin patterns alone were sufficient to arrest the migration of CAFs and defects within the CAFs layer, even if CAFs are unable to produce fibronectin.

### A minimal model of active nematic with coupling to a dynamic, passive nematic field recapitulates the observed aging and freezing dynamics

So far, experimental observations show that CAFs behave as an active nematic layer that produces a fibronectin matrix with coinciding nematic order and that CAFs respond and align with a pre-existing nematically order matrix; as opposed to active nematic systems described so far, which exhibit a dynamic stationary state, this coupling was found to induce aging dynamics that eventually lead to a frozen state with immobile topological defects and no observable cellular flow despite local active fluctuations. In order to show that indeed, with minimal hypotheses, the coupling of an active nematic cell monolayer with a nematic ECM that it deposits generically can lead to the slow down and eventually freezing of both defect dynamics and cellular flows, we developed a minimal theoretical model that recapitulates our experimental observations (**Fig. 5; Supp. Fig. 9**). The model describes minimally the dynamics of the nematic order parameter *Q* of the CAF layer building up on classical active nematohydrodynamics ^17, 19, 26, 29^ (see supplementary model (SM) for details):

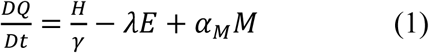

where 𝐷 𝑄⁄𝐷𝑡 = 𝜕_𝑡_𝑄 + (𝑣. 𝛻)𝑄 − (𝑄. 𝛺 − 𝛺. 𝑄) is the corotational convective derivative, 𝐻 = −𝛿 𝐹⁄𝛿 𝑄 is the so-called molecular field, 𝛾 the rotational viscosity, 𝛺 = (𝛻𝑣 − (𝛻𝑣)^𝑇^)⁄2 the vorticity, 𝜆 a flow alignment coupling, 𝐸 = (𝛻𝑣 + (𝛻𝑣)^𝑇^)⁄2 the strain rate tensor, and 𝐹 the standard free energy of a nematic field that would control the dynamics in the absence of activity (see SM). The key ingredient that we introduce is the nematic matrix field 𝑀 that encodes the strength and direction of the fibronectin matrix alignment; in turn, the response of the CAF nematic order to the matrix is taken into account *via* a phenomenological active coupling 𝛼_𝑀_ in (1), which favors the alignment of *Q* with *M*. The assembly-degradation dynamics of 𝑀 can be minimally encoded as

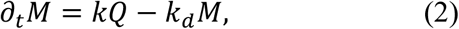

where *k* is the fibronectin deposition rate (which can be *M* dependent to ensure that *M* remains finite) and 𝑘_𝑑_ a degradation rate (assumed vanishing following our experimental observations). Note that a comparable coupling between cell orientation and matrix organization has been proposed in the literature on the basis of agent based models, designed to analyze heterogeneous cell assemblies ^37, 38^; instead, we focus here on confluent cell monolayers for which our hydrodynamic description is better suited. Finally, force balance on the CAF’s layer can be written

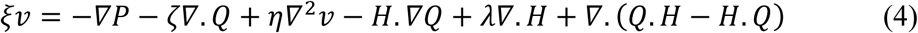

where 𝜉(𝑀) is a potentially anisotropic friction tensor that can depend on 𝑀, *P* a pressure, 𝜁 a classical phenomenological coupling defining the active stress, and 𝜂 the viscosity. For the sake of simplicity, we first assumed that the friction 𝜉 is independent of 𝑀, and considered limit regimes of either fully compressible (neglecting *P* in (4)) or incompressible (where *P* is a Lagrange multiplier enforcing 𝛻. 𝑣 = 0). Numerical analysis of the model, in agreement with analytical arguments following Ref ^39^, shows that the velocity of isolated +1/2, comet defects is given in steady state by

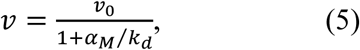

where 𝑣_0_ is a velocity scale independent of 𝑘_𝑑_ (see SM), and can thus be dramatically decreased for either a slow matrix degradation rate (𝑘_𝑑_ → 0) or strong alignment interaction 𝛼_𝑀_ (**Fig. 5B**), which can be interpreted as the relaxation rate of the nematic field *Q* to the matrix field. The slowdown and even arrest of defects, as is observed in experiments, can thus be triggered by the alignment interaction 𝛼_𝑀_ only, even for a friction 𝜉 assumed independent of 𝑀. The numerical analysis of the model in general cases with multiple defects shows that this sole process reproduces the expected difference in the defect velocities for siCtrl and siFN (**Fig. 5D-F; Supp. Fig. 9D; Movie S1**)𝑘_𝑑_ → 0f note, in this case, cellular flows are maintained even in the limit (￼) of arrested defects (see SM). To reproduce the observed decrease of both defects speed and cellular flow speeds in siCtrl experiments, we thus made the further hypothesis that friction is anisotropic and depends on *M* acc𝜉 = *aI* + *bM*rding to ￼𝑎, 𝑏where ￼ are phenomenological functions of the norm of *M* given in SM. This form is consistent with the observed anisotropy in single-cell dynamics (**Fig. 2B**), and indeed yields an aging dynamics of the flow field, with a significant decrease in cellular flow speeds over time (**Fig. 5C-E; Supp. Fig. 9E; Movie S1**). This slowdown is, at long times, controlled by the matrix deposition rate *k* is found to be spatially uniform (and thus scale independent, see **Supp. Fig. 9E, F**), in agreement with experiments. A further confirmation of the model is given by the experiments depicted in Fig. 5, where CAFs are plated on a pre-existing FN pattern. This can be readily implemented in the model by assuming a given random initial condition for the matrix field *M*. We find𝛼_𝑀_by solving numerically the model ￼the dynamics of the cellular field 𝛼_𝑀_ in this case quickly relaxes (within a time scale controlled by 1/￼) to a frozen state with *Q* matching *M*, in agreement with experiments (**Supp. Fig. 9G-H**). Finally, the model shows, with minimal hypotheses, that the coupling of an active nematic cell monolayer with a nematic ECM that it deposits generically leads to the slowdown and eventually freezing of both defect dynamics and cellular flows, and this is spatially uniform.

**Figure 5.**
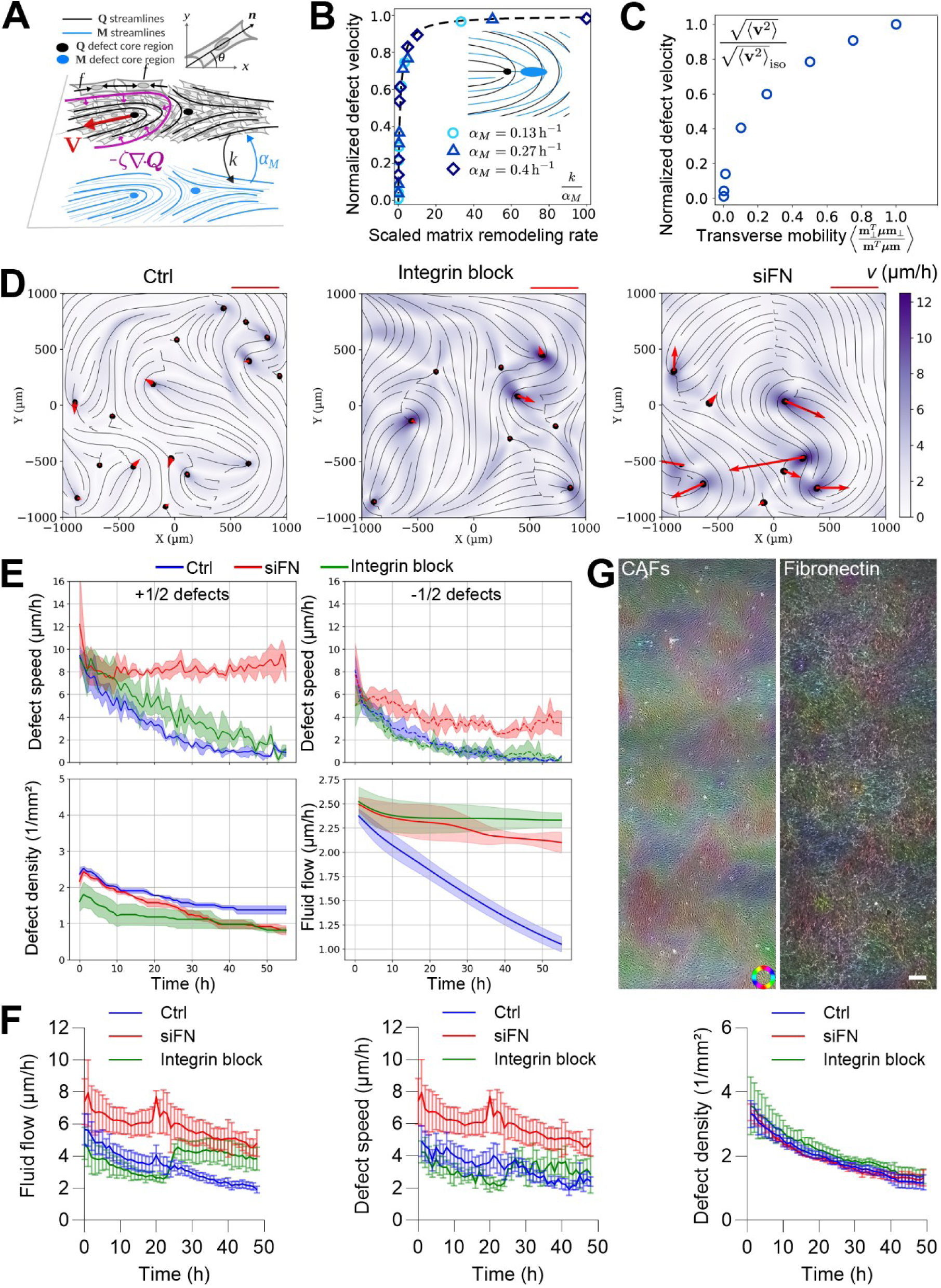
Theoretical model. A) The fibroblast layer is modeled as an active nematic. A vector, n, along the major axis of the cells is used to define the nematic order parameter, Q. The fibroblasts exert an active stress, -ζQ, where the parameter, ζ, is introduced in Eq. 4, and ζ < 0 for the contractile fibroblast layer. This stress originates from single-cell force compressive dipoles, denoted as f. Fibroblasts deposit fibers with nematic alignment (M) onto a surface at a rate of k. They also actively reorient to the matrix with coupling αM. B) The velocity of the comoving defects as a function of the scaled remodeling rate, k/α_M, for different values of matrix coupling, αM. The velocities are normalized by the numerical velocity value as k→+∞. The theoretical scaling for v/v₀ predicted in Eq. (5) is indicated by the black dashed line. A numerical visualization of interacting fibroblast (blue) and matrix (black) defects is added. The core region is shown as the region for which S < 0.42 and SM < 0.35. C) The mean absolute value of the fluid flow velocity on day 3 as a function of friction anisotropy in a control case. The average velocity tends to zero when mobility friction goes to zero. D) A comparison of siCtrl, siFN, and integrin block (β1 integrin blocking antibody, AIIB2) scenarios on day 3. The color map corresponds to fluid flow velocity (FV), and the arrows correspond to defect velocities. The velocity measure bar corresponds to 5 µm/h (DV). siCtrl displays low FV and DV; integrin block displays low DV and high FV; and siFN displays both high FV and DV. See also **Movie S1**. E) Top: Defect velocity for +½ (left) and -½ (right) defects in different scenarios: siCtrl (blue line), siFN (red line), and integrin block (β1 integrin blocking antibody, AIIB2, green line). Bottom: Defect density (left) and mean fluid flow velocity (right) in different scenarios: siCtrl (blue line), siFN (red line), and integrin block (β1 integrin blocking antibody, AIIB2, green line). Due to full defect freezing, siCtrl has a higher defect density. Conversely, siFN defects have higher velocities. Integrin block and siCtrl share similar mean fluid flow velocity. F) Evolution of the mean of the velocity field (left), the defect speed (middle) and the defect density (right) of CAF layers in control (combined results of siCtrl and DMSO; Ctrl, bleu line), siFN (red line), or integrin block (β1 integrin blocking antibody, AIIB2, green line) scenarios. AIIB2 was added 24h after the start of the time-lapse. The lines represent the mean of >3 independent experiments, and the error bars represent the standard error of the mean. I) Orientation of CAFs based on bright field (left) and orientation of fibronectin network (right) 24 h after addition of a β1 integrin blocking antibody (AIIB2). Colored circle: orientation colormap. Scale bar: 200 µm.

To further challenge the model and get insights into the molecular mechanisms involved in the CAFs / fibronectin matrix interactions, we used a β1 integrin-blocking antibody (AIIB2). Remarkably, shortly after integrin inhibition (induced after 24h of unperturbed dynamics), we observed a clear reawakening of the CAFs’ dynamics, characterized by increased cellular flows, while importantly, the defect density and velocity remained comparable to the control (**Fig. 5F**). Strikingly, immunofluorescence staining conducted at the end of 48 hours showed that the nematic organization of fibronectin remained intact (**Fig. 5G**, correlation of 0.94+/- 0.01, 3 independent experiments, between CAFs and fibronectin layer), suggesting that the aligning interaction of cells with the matrix is preserved. This suggests that integrins indeed mediate a coupling between cellular and ECM nematic fields and are responsible for the slowdown of cellular flows observed in untreated monolayers. However, this interaction primarily affects the effective friction of the cell monolayer with the matrix (which indeed is expected to slow down the cell dynamics, see ^17, 40^), while preserving the local alignment of cells with the matrix. The model successfully recapitulates these observations, as it indeed shows that the orientational coupling to the matrix alone (𝛼_𝑀_) is sufficient to freeze the dynamics of the defects, with, however, non-vanishing cell flows, while a complete freezing of cell flows requires, in addition, an increased friction 𝜉 (**Fig. 5C, D; SM**).

Altogether, an active hydrodynamic model of the CAF layer shows that the interaction with a deposited nematic matrix induces feedback that can lead to a slowdown and eventually freezing of the dynamics of both defects and cellular flows.

### Fibronectin loss–induced fluidization of the CAF capsule enables basement-membrane breach

In tumors, the spatial organization of CAFs and the ECM is tightly linked to the stage of invasion and is described as tumor-associated collagen signatures (TACS) ^8, 41, 42^. In non-invasive regions (TACS-2), CAFs and ECM align parallel to the tumor edge, forming a capsule-like structure, well documented in mouse tumors. To assess whether such a capsule also exists in human cancers, we analyzed freshly resected colorectal tumor tissue. The tumor was rapidly processed—fixed, sectioned into 700 µm slices, stained, optically cleared, and imaged using two-photon confocal microscopy. This revealed a distinct CAF capsule in human tumors, closely resembling the one described in mouse models (**Fig. 6A**).

**Figure 6.**
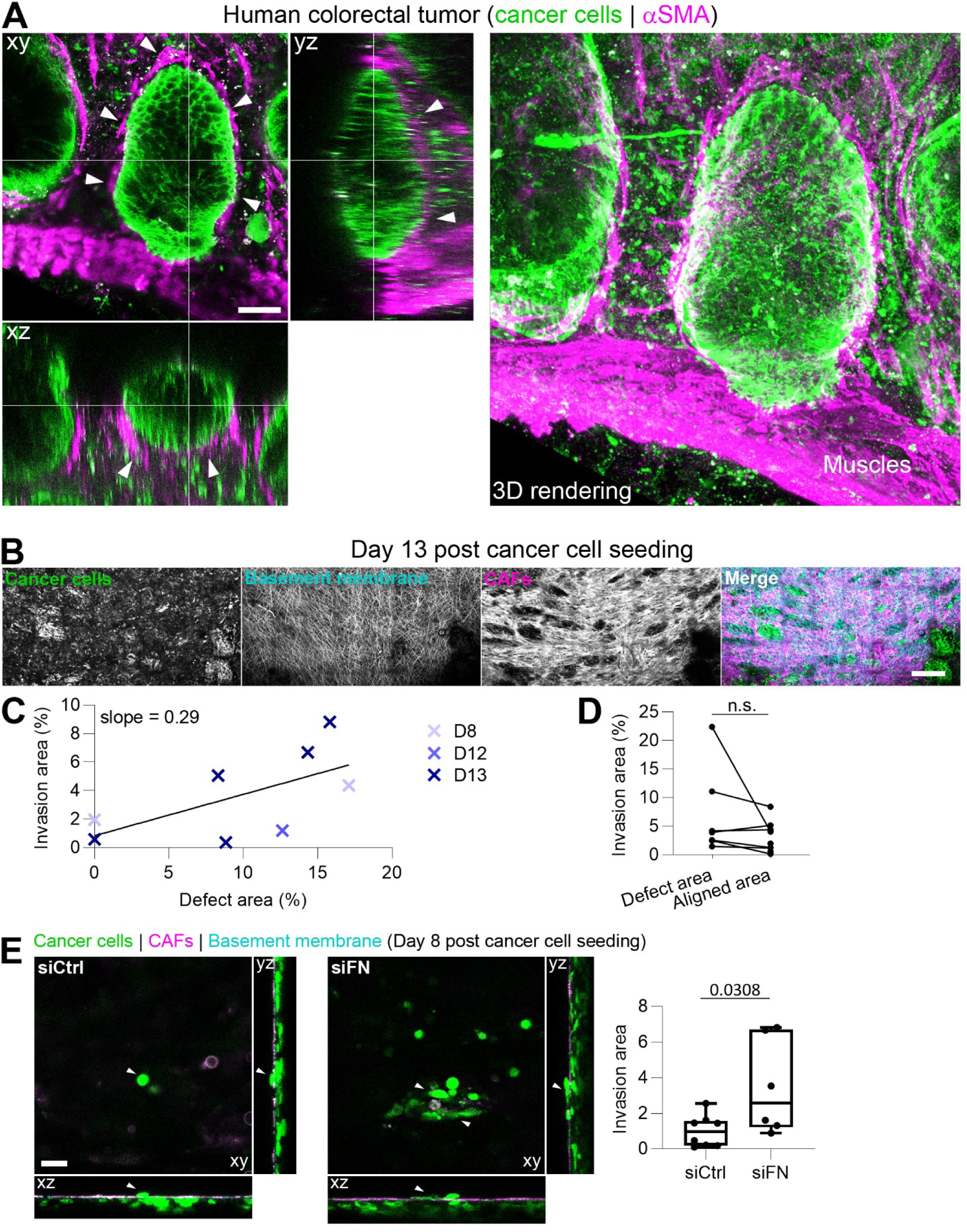
Loss of fibronectin in CAFs enhances cancer cell invasion. A) 3D imaging of a human colorectal tumor showing cancer cells (EpCAM, green) and CAFs (aSMA, magenta). Note the organization of CAFs around the group of cancer cells (white arrowheads). Orthogonal view and 3D images are shown. Scale bar: 50 µm. B) 3D imaging of HCT116 cancer cells (green), CAFs (magenta) on a native basement membrane (cyan), 13 days post cancer cell seeding. Maximum projection images are shown. Scale bar: 500 μm. C) Correlation between cancer cell invasion area (% of the FOV) and defect area (% of the FOV) inside the CAFs layer. Each cross represents one invasion assay, with the time of invasion indicated in days. D) Proportion of cancer cells invasion (% of the FOV) at defect areas and aligned areas. One pair of dots represents one invasion assay. Boxplot: middle bar= median, edges bars= 25th and 757th percentiles, whiskers= extent of data. Paired two-sided t-test, p-value = 0.2357. E) 3D imaging of HCT116 cancer cells (green), CAFs (magenta) on a native basement membrane (cyan), 8 days post cancer cell seeding. CAFs were transfected with either control siRNA (siCtrl) or siRNA against fibronectin (siFN). The xy plane is chosen so that cancer cells that invaded below the CAF layer are visible (see white arrowheads). Scale bar: 100 μm. Right: invasion area (relative to siCtrl) in an invasion assay where CAFs were transfected with either control siRNA (siCtrl) or siRNA against fibronectin (siFN). One dot represents one invasion assay. Boxplot: middle bar= median, edges bars= 25th and 757th percentiles, whiskers= extent of data. Two-sided unpaired t-test.

In invasive regions (TACS-3), CAFs and ECM reorient perpendicularly, creating tracks that facilitate tumor dissemination ^8, 41, 42^. This suggests that defects in the CAF capsule could serve as sites where CAFs reorient, driving the transition from TACS-2 to TACS-3. Alternatively, mechanical stresses at defects may “pinch” the underlying substrate and tumor cells, piercing the CAF layer and enabling invasion.

To directly test whether defects in the CAF capsule promote cancer-cell invasion, we used a native basement-membrane invasion assay ^43^. CAFs were plated on one side of decellularized mouse mesenteries and allowed to establish nematic order. Cancer cells were then seeded on the opposite side. Cultures were maintained for up to 13 days, and invasion events were quantified at CAF-layer defects versus aligned regions (**Fig. 6B**). Invasion increased over time, with a few events on day 8 and a rise by day 13. Although there was a weak correlation between the number of defects and invasion (**Fig. 6C**), cancer cells did not invade preferentially at defect sites compared to well-aligned areas (**Fig. 6D**). However, invasion itself perturbed the CAF layer, potentially disrupting nematic order and complicating interpretation.

Strikingly, depletion of fibronectin in CAFs markedly enhanced invasion. Because fibronectin stabilizes the “frozen” nematic state of the CAF capsule, its loss increases nematic fluidity and weakens barrier function, permitting greater cancer-cell passage (**Fig. 6E**).

Together, these results demonstrate that CAF-layer integrity is crucial for restraining tumor invasion. While defects in the capsule can contribute, it is the transition from a stable, fibronectin-stiffened (“frozen”) nematic architecture to a more fluid state that critically undermines the barrier and enables cancer cells to invade.

## Discussion

Our study identifies a self-organized feedback between CAFs and fibronectin that transforms the CAF capsule into a frozen active nematic. CAFs align nematically and deposit fibronectin fibers with nearly perfect orientational correlation. In turn, this fibronectin matrix stabilizes CAF alignment, increases frictional coupling, and progressively arrests defect motion and cellular flows—yielding an “active nematic solid” where defects are immobilized but remain sites of concentrated mechanical stress. Perturbation experiments confirmed fibronectin’s central role: its depletion restored CAF and defect motility, while pre-patterned fibronectin nematics alone were sufficient to arrest CAF dynamics. A minimal active–passive nematic model reproduced this aging and freezing behavior, and integrin blockade experiments demonstrated that frictional pinning to fibronectin is key to flow arrest.

It is important to emphasize how unique this ECM-driven nematic arrest is compared to other active nematic systems. In classic active nematics, the active stresses continuously generate ±1/2 defect pairs. The +1/2 defects are motile, leading to perpetual defect chaos (active turbulence) with a steady-state defect density. Only by applying external constraints can such systems achieve long-lived orientational order or pinned defects. For instance, earlier studies have shown that imposing substrate anisotropy or micropatterned topography can orient cells globally and even stabilize defects in place. However, these approaches rely on extrinsic cues or boundary conditions to control the active nematic. In our system, by contrast, the alignment and arrest emerge autonomously via the cell–matrix feedback: the cells themselves generate the aligning field (fibronectin fibers) and the increased friction, without any engineered pattern in the substrate. Thus, there is an intrinsic suppression of active turbulence in a living cell monolayer through a two-way coupling with a secreted matrix. This result enriches the understanding of active matter by showing that active nematics can transition into a new regime when long-lived passive structures are co-generated by the system. Conceptually, the fibronectin matrix here plays the role of an internal field that stores the memory of the alignment and builds up over time. Aging and slow-down phenomena are well-known in passive soft glassy materials; our findings show an active analog where the agents (cells) themselves produce a structure that causes their collective dynamics to vitrify. We also verified that this behavior is not limited to CAFs from tumors: normal fibroblasts (NAFs) exhibited the same ability to form an aligned fibronectin network and reach a frozen nematic state. This suggests a general biophysical mechanism whereby connective tissue cells can self-organize to suppress large-scale motion – potentially relevant in wound healing, fibrosis, and other contexts where fibroblasts densify and lay down matrix.

Functionally, the CAF capsule forms a barrier that restrains tumor spread. While defects correlate with increased invasion, they do not represent preferred breach points; instead, the decisive step occurs when the capsule fluidizes due to fibronectin loss. Invasion assays showed that fibronectin depletion markedly enhanced cancer-cell invasion, underscoring that fibronectin-mediated nematic freezing is essential to barrier integrity. Thus, defects are permissive but capsule fluidization is decisive for invasion. More broadly, our results demonstrate that active matter systems can arrest their own dynamics through ECM feedback, revealing a biophysical mechanism by which stromal organization governs tumor containment and progression.

## Acknowledgments

We thank all members of the DMV lab (especially Réda Bouras for his assistance with gel generation and cell culture) and Silberzan lab for helpful discussions. The authors greatly acknowledge the Cell and Tissue Imaging (PICT-IBiSA), Institut Curie, member of the national infrastructure France-BioImaging (https://ror.org/01y7vt929) supported by the French National Research Agency (ANR-24-INBS-0005 FBI BIOGEN). This work was supported by the Fondation pour la Recherche Médicale (FRM N° DGE20111123020), the Cancerople-IdF (n°2012-2-EML-04-IC-1), InCA (Cancer National Institute, n° 2011-1-LABEL-IC-4). This work received funding from the European Union’s Horizon 2020 research and innovation program: European Research Council (ERC) under the grant agreement CoG 772487 (DMV), AdvG 883739 (XT), AvdG 101019835 (BL), SyG SHAPINCELLFATE (RV), Aviesan ITMO Cancer « Convention Frontières du Vivant» N°18CF156-00 (CJ), Fondation pour la Recherche Médicale - FRM FDT202106013007 (CJ), Institut National du Cancer INCa 16712 (BL), the Ligue Contre le Cancer (Equipe labellisée 2019) (BL), LABEX Who Am I? (ANR-11-LABX-0071) (BL), Fondation ARC pour la Recherche sur le Cancer (CPG), et ANR (RV), EPSRC grant no EP/R014604/1 (AM). AM was supported in part by a TALENT fellowship awarded by CY Cergy Paris Université. AM would like to thank the Isaac Newton Institute for Mathematical Sciences, Cambridge, for support and hospitality during the program New statistical physics in living matter: non-equilibrium states under adaptive control, where a part of the work on this paper was undertaken.

## Supplementary material

**Movie S1.** Evolution of nematic order and cell dynamics for Ctrl (left), siFN (middle) and integrin block (right) conditions. Orientational streamlines represent the nematic alignment of fibroblasts, with topological defects marked as black dots. The color map shows the velocity field (µm/h). Movies start 10 h after the beginning of the simulation, which is performed in a closed system of size 2250 × 2250 µm². The red scale bar represents a length of 400 µm. **Supplementary model (SM).**

## Material and methods

### Human sample

The tissue was fixed in 4% PFA (Electron Microscopy Sciences cat. #15711) for 1h at RT, washed 3 times in PBS, and then embedded in 3% agarose (Invitrogen cat. #16520050). 700 μm-thick sections were generated using a vibratome (Leica) with a 0.4 mm/s speed and 2 mm amplitude. About 2 to 3 sections were placed flat inside a cryomold, and after permeabilizing with 2% Triton X-100 in PBS for 3 days at RT, tissue sections were incubated in blocking buffer [10% fetal bovine serum, 1% Triton X-100, 4% DMSO in PBS] for 1 day at RT. Primary antibodies were incubated for 2 days at RT (mouse anti-aSMA Sigma-Aldrich cat. #A2547, rabbit anti-EpCAM Abcam cat. #ab71976). Samples were rinsed with 0.2% Tween and incubated with secondary antibodies for 1 day at RT (goat anti-mouse Alexa Fluor 488 and goat anti-rabbit Alexa Fluor 546, Invitrogen). All incubation steps described previously were done under gentle orbital shaking conditions. To mount such thick tissues, spacers were placed on a microscopy slide, and the resulting chamber was filled with clearing solution (RapiClear; cat. #RC149001). The tissues were immersed in the clearing solution, and a coverslip was mounted to seal the chamber. Samples were imaged on a 2-photon microscope (Leica SP8 NLO) with a 25X objective (HC IRAPO L 25x/1.0 W, ref # 15506340).

### Cell culture

For this study, we have used CAFs isolated from colon tumor samples of a patient treated at Institut Curie Hospital, Paris, with the patient’s written consent and approval of the local ethics committee. CAFs were extracted from fresh tumor samples and immortalized as described earlier ^1–3^. CAFs were cultured on 30 kPa soft substrate plates (ExCellness – PrimeCoat, Biotech SA). Before being used, plates were plasma-treated for 30 s at 800 mTorr and then coated with 5 µg/mL of monomeric collagen in DMEM supplemented with 2X Anti-Anti, 12.5 µg/mL Metronidazole B (Braun), and 4 µg/mL Ciprofloxacin (Panpharma). After incubation overnight at 37°C, 5% CO_2,_ dishes were washed with medium before adding the CAFs medium (DMEM 5% FBS and 1X ITS-A) and seeding approximately 1×10^6^ CAFs. CAFs were left to grow for one week at 37°C 5% CO_2_ before being used for experiments.

The colon cancer cell line HCT116 expressing cytoplasmic GFP was cultured in DMEM supplemented with 5% FBS, as previously described ^3^.

### Basement membrane preparation

Basement membranes were extracted from the mesentery of B6N mice females between 8 to 9 months of age (purchased from Charles River). The gut was dissected, and the mesentery was gently extracted using tweezers, with frequent hydration using PBS supplemented with 2X Anti-Anti. A piece of mesentery was secured in between two magnetic rings (made in house and put in PBS containing 2X Anti-Anti. About six mesentery pieces could be extracted from one mouse. Mesenteries were then decellularized by incubating in 1M NH4OH for 40 min at room temperature. Decellularized mesenteries were then washed for 40 min at room temperature in PBS supplemented with 2X Anti-Anti. Sulfo-sampah was applied on both sides of the mesentery and activated using a UV lamp for 5 min. Mesenteries were then washed for 3 min with 10 mM HEPES and two times for 3 min with PBS supplemented with 2X Anti-Anti while shaking. Finally, mesenteries were incubated at 4°C overnight with 100 μg/mL of monomeric collagen I in PBS supplemented with 2X Anti-Anti.

### Invasion assay

Invasion assays were performed on mesenteries mounted on magnetic rings and were cultivated in DMEM with either 10% or 1% FBS, 1X ITS-A, 2X Anti-Anti, 12.5 μg/mL Metronidazole (B. Braun), and 4 μg/mL Ciprofloxacin (Panpharma). First, mesenteries on rings were placed in 12-well plates with 700 μL medium to have the bottom part of the basement membrane immersed in media. 2×10^5^ CAFs were seeded on the top part of the mesentery in 200 μL medium per ring. Rings were carefully transferred to the incubator at 37°C and 5% CO2. After 40 min, 10% FBS medium was added into each well up to 2 mL. Two days later, rings with mesenteries were turned upside down, and 2×10^5^ HCT116 cells expressing GFP were added on the opposite side of the CAFs following the same process as for the fibroblasts. The day after, the medium was changed to reduce the FBS concentration from 10% to 1%. The construct was then incubated at 37°C and 5% CO2, with half of the medium being changed every two days until the desired invasion time point. Mesenteries were then fixed with 4% PFA for 20 min at room temperature and imaged using an inverted laser scanning confocal Leica SP8 coupled to a multi-photon femtosecond Chameleon Vision II laser (680-1350 nm; Coherent) and a 25X water immersion lens (NA, 1.0). The basement membrane was imaged using second harmonic generation.

### Invasion assay analysis

Matlab and Fiji were used to do the analysis.

#### Correlation defect/invasion

For each experiment, the defect area and cancer invasion masks were manually drawn using Fiji. The masks obtained were transformed into 1 and 0 matrices. The no-defect areas mask was obtained by reversing the 0 and the 1 of the defect area mask. The cancer invasion inside the defect area mask was obtained by multiplying the cancer invasion mask and the defect area mask. The cancer invasion outside the defect area was obtained by multiplying the cancer invasion mask and the no-defect area mask. For each mask, the size of the positive area was calculated. The field of view area, called the total area, is also calculated. Values of interest were then calculated as percentages. Then, the proportion of cancer invasion area toward the proportion of defect area, and the fraction of cancer invasion in the defect area toward the proportion of cancer invasion area were plotted. The proportion of cancer invasion in defect and no-defect areas were pulled together into boxplots for each condition.

#### Invasion siCtrl/siFN

Using a custom-made Matlab code, z-stacks were flattened based on the basement membrane channel. Then, using the *Orthogonal view* tool in Fiji, the planes below the basement membrane were identified, and the total area of cancer cells present in these planes were measured.

### LifeAct GFP and GFP transfection

Fibroblasts were transfected using lentiviral infection. Lentiviruses containing the LifeAct GFP or the GFP (Addgene; cat. #65656) plasmid with 4 µg/mL of Polybrene were added to fibroblasts and incubated overnight at 37°C, 5% CO_2_. After incubation, fibroblasts were washed several times with media and then incubated at 37°C 5% CO_2_. After a few days, the transfection success was checked using a fluorescent microscope. Once the lifeAct-GFP fibroblast population reached 15 to 20 million cells, fibroblasts were sorted to remove GFP-negative fibroblasts. Sorted fibroblasts were then cultured as described above.

### PAA gel preparation

PAA gels were prepared on Glass-bottom dishes (Fluorodish, plate 35 mm Ø, Glass 23 mm Ø, WPI) as previously described ^44^. First, plates were treated with 3-amino-propyltrimethoxysilane (diluted 1:2 with PBS; Sigma-Aldrich/Merck) for 15 min at RT. After three washes with MilliQ water and air drying, dishes were treated with glutaraldehyde 0.25% in PBS (Sigma-Aldrich/Merck) for 30 min at RT. After three washes with MilliQ water and air drying, PAA solution for gels was prepared (see Table 1). Alternatively, a treatment with Bind-silane (Sigma-Aldrich) dissolved in absolute ethanol (PanReac) and acetic acid (Sigma-Aldrich) at volume proportions of 1:12:1 for 10 min at RT was done. After three washes with absolute ethanol and air drying, PAA solution for gels was prepared (see Table 1). For traction force microscopy experiments, 5 µL of 0.2 µm red beads FluoSpheres carboxylate-modified (580/605, Invitrogen) were added to the PAA gel solution. 22.5 µL drops of the PAA gel solution were put in the middle of the dish and covered with 18 mm diameter glass coverslips. PAA gels were let to polymerize for one hour at RT. Once PAA was polymerized, PBS was poured into dishes, and glass coverslips were removed using a scalpel and a tweezer. PBS was removed, and 100 µL of 2 mg/mL sulfo-sanpah (Sigma-Aldrich/Merck) was added only onto PAA gels. Dishes with sulfo-sanpah were then treated with ultraviolet light (365 nm; 5 cm from source) for 5 min. PAA gels were washed briefly with 10 mM HEPES, followed by two washes with PBS. Dishes were then incubated with 200 µL of 100 µg/mL monomeric collagen I (Rat tail origin, Corning) diluted in 0.2% acetic acid, or 100 µg/mL laminin-111 (Thermo Fisher Scientific cat. #23017015), pipetted only on the PAA gel at 4°C overnight. Dishes were washed with PBS before the addition of CAFs and culture medium.

**Table 1.** Volumes (in µl) of reagents to prepare different stiffness PAA gels.

| Stiffness (Young Modulus) | PBS | Acrylamide 40% | Bis-acrylamide 2% | APS | TEMED |
| --- | --- | --- | --- | --- | --- |
| 5 kPa | 382,95 | 93,3 | 11 | 2,5 | 0,25 |
| 11 kPa | 368,5 | 93,75 | 25 | 2,5 | 0,25 |
| 30 kPa | 299,75 | 150 | 37,5 | 2,5 | 0,25 |

### Time-lapse imaging of nematic ordering evolution over time

15 x 10^4^ CAFs treated with either control siRNA (siCtrl) or siRNA against fibronectin (siFN), were seeded on 11 kPa PAA gels. Alternatively, 15 x 10^4^ CAFs were seeded on 5, 11, or 30 kPa PAA gels. After plating, fibroblasts were incubated in DMEM medium containing 10% FBS, 1X ITS-A, 2X Anti-Anti, 12.5 µg/mL Metronidazole (B. Braun), and 4 µg/mL Ciprofloxacin (Panpharma) at 37°C 5% CO_2_ for 24 h before imaging. CAFs were imaged using an inverted Eclipse Ti-2 microscope (Nikon) driven by NIS elements (Nikon) with a fully motorized stage, a 10X (NA, 0.3) objective, and an incubation system at 37°C, 5% CO_2_. The entire CAFs layers of each condition were imaged for 50 h, with images taken every 1h.

### 2D traction force microscopy experiment

CAFs were seeded on 11 kPa PAA gel with red beads (580/605, Invitrogen, F8810) and cultured in DMEM medium containing 10% FBS, 1X ITS-A, 2X Anti-Anti, 12.5 µg/mL Metronidazole (B. Braun), and 4 µg/mL Ciprofloxacin (Panpharma) at 37°C 5% CO_2_. The number of cells seeded depended on the desired cell density at the moment of imaging. During siRNA experiment, CAFs were treated with either control siRNA (siCtrl) or siRNA against fibronectin (siFN). CAFs were imaged using an inverted Eclipse Ti-E microscope (NIKON) driven by Metamorph software (v.7.8.13.0) with a fully motorized stage, a 10X (NA, 0.3) objective, and an incubation system at 37°C, 5% CO_2_. Beads were imaged using 562/40 nm excitation and Bright Field was used to image the CAFs. For some experiments, 472/30 nm excitation was used to image LifeAct-GFP CAFs, while 377/50 nm excitation was used for Hoechst staining. For each region of interest focusing on one type of topology: two single z slices were taken, one focusing on the fibroblast plane and one focusing on the beads. Time-lapse imaging lasted for ∼16h, with images taken every 15 min. At the end of imaging, dishes kept on the microscope stage were carefully washed with PBS before cells were removed with TrypLE Express (ThermoFisher). Once cells detached from PAA gels, one-time point acquisition was performed for all positions to obtain the reference points for relaxed gel.

### 2D traction analysis

Using Matlab, traction force experiments were analyzed using scripts previously developed ^45^. First, data were separated into folders corresponding to each position, then positions were treated separately. For each position, all images were corrected for potential drift, aligned, and cropped the same way using the image of beads after trypsin treatment as a reference. Using the Particle Image Velocity (PIV) method, the displacement of the beads between traction and trypsin images was measured on eight-by-eight pixels squares along all images with overlapping between each square to precise the measure. From the displacement and the gel settings (finite thickness), traction forces were calculated using Fourier-transform traction microscopy. The traction magnitudes’ maps were plotted for each time point and traction vectors were plotted on top of the cell images.

### Internal stress analysis

Internal stresses of the cell layer were analyzed using Bayesian Inversion Stress Microscopy (BISM) script previously developed ^46^ using Matlab. Internal stresses were calculated from the traction forces obtained previously using Bayesian inversion theory. BISM doesn’t depend on the physical property of the system such as tissue rheology, and thus doesn’t need boundary conditions. However, the system needs to have a small height compared to its planar surface and is not reliable very close to the boundary. BISM is based on strong statistical hypotheses such as the Gaussian distribution of the stresses. Tests performed by Vicent Nier have shown that BISM is robust and could be applied to different systems as long as they validate the height condition. Only defects inside the layer were taken to avoid boundary issues. In addition, the height condition was validated in this study as the fields of view are 825.6 µm by 598.56 µm, and CAFs height is about 10-15 µm, so BISM can be applied.

### Defects averaging

Matlab was used to do the analysis. For each experiment, positions were separated into different types of topology: comet defects, triangle defects, and aligned areas. Positions of each type of topology were then aligned in the same direction. To do so, the core and direction places of the defect/aligned area were pointed out of the bright field images (see Fig. 2C for a visual representation of core and direction places for each type of topology). For each position and time point, core-centered circles with a radius of 800 pixels were used as masks. These circular masks were applied to traction force, and internal stress maps to only keep data within the masks. Rotations of these circular masks with traction force and internal stresses were then performed to align their directions. Finally, for each type of topology, the averages of all the circles were calculated to obtain their specific force pattern organization.

### Fibronectin and N-cadherin localization over time

CAFs mixed with 2% GFP-expressing CAFs were seeded, with a total of 2 x 10^4^ cells, on 11 kPa PAA gels. Cells were fixed for 10 min at RT in 4% PFA, 3 or 8 days post-seeding. After several PBS washes, cells were permeabilized with 0.5% Triton X-100 in PBS for 15 min at RT. Cells were then blocked for 45 min at RT in blocking solution (10% FBS, 0.05% Triton X-100 in PBS). Primary antibodies (see Table 2) were incubated in blocking solution for 1 h at RT. Following several washes in 0.05% Tween-20 in PBS wash solution, cells were incubated with secondary antibodies and Phalloidin (see Table 2) for 1 h at RT. Cells were then washed multiple times with wash solution and stored in PBS at 4°C until imaging. Cells were imaged using an inverted Eclipse Ti-E microscope (Nikon) with a spinning disk CSU-W1 (Yokogawa) integrated into Metamorph software (v.7.10.2.240) by Gataca Systems, utilizing a 40X oil immersion lens (NA, 1.3).

**Table 2.**
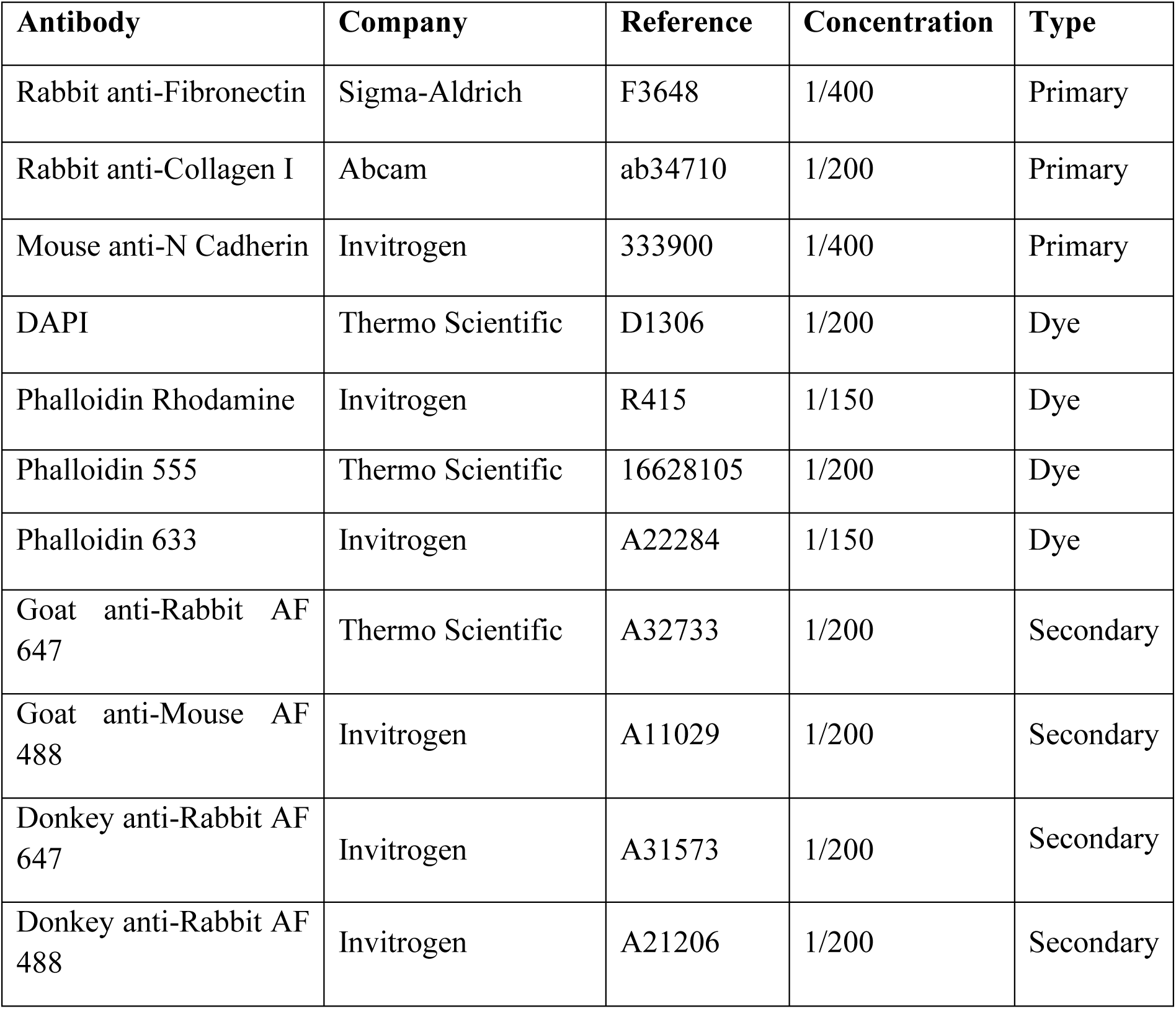
The list of antibodies used in this study.

N-Cadherin and fibronectin intensities at cell edges were quantified using a semi-automatic custom macro developed in Fiji software. For each image stack, a representative middle slice was selected.

#### N-cadherin

GFP+ cells were segmented as particles larger than 600 µm². for each segmented GFP+ cell ROI, a 3-pixel-wide polyline selection was generated, corresponding to the outline of the cell ROI (*i.e.* membrane signal). Next, the cell ROI was eroded by 2 pixels, and a second 3-pixel-wide polyline selection was generated, corresponding to the outline of the eroded cell ROI (*i.e.* cytoplasmic signal). Mean pixel intensities were measured for both lines. The mean intensity of the cytoplasm line was subtracted from that of the membrane line to correct for background. Values from day 8 were normalized to those from day 3. Manual adjustments were performed as necessary to specifically measure cell-cell contact borders.

#### Fibronectin

for each GFP+ cell, a 100-pixel-wide line was manually drawn across the cytoplasm and positioned at the level of the nucleus. The line was positioned perpendicular to the long axis to cover the entire cell width. From this line, a fluorescence intensity plot profile was generated, representing fibronectin signal across the cell width. To allow comparison between cells of different sizes, the cell width was normalized in percentage. Each profile was divided into bins, and the average fluorescence intensity was calculated within each bin (*e.g.*, 0–5%, 5–10%, etc.). Final data represent the mean intensity distribution across normalized widths for all analyzed cells.

### Immunostaining

Immunostainings were performed using CAFs growing on 11 kPa PAA gels described above. Cells were pre-extracted before being fixed. To do so, a 50 mL 2X PEM solution was prepared in MilliQ water with 3 g PiPES, 1 M MgCl_2_, 100 mM EGTA and pH adjusted at 6.9 using KOH. First, cells were washed with PBS and then incubated for 30 s at RT with the extraction solution (0.2% Triton X-100, 50% 2X PEM, 4% PEG 35000 and 5 µM Phalloidin in PBS). Cells were washed two times with the rinse solution (50% 2X PEM, 2 µM Phalloidin in MilliQ water) and then incubated in 4% PFA in PBS solution for 20 min at RT. After several washes with PBS, cells were incubated with the primary antibodies for 1h at RT. Cells were washed several times with PBS and then incubated with the secondary antibodies for 1h at RT. Cells were washed several times with PBS and mounted on glass slides with AquaPolyMount. Stained cells were kept at 4°C before imaging. All the antibodies used during this study and their concentration are listed in Table 2. Cells were imaged using either an inverted Eclipse Ti-E microscope (NIKON) with spinning disk CSU-W1 (Yokogawa) integrated into Metamorph software (v.7.10.2.240) by Gataca Systems, a 60X water immersion lens (NA, 1.27), a 40X water immersion lens (NA, 1.15), and a 20X (NA, 0.45) objective; or an inverted laser scanning confocal Leica SP8 coupled to a multi-photon femtosecond Chameleon Vision II laser (680-1350 nm; Coherent) and a 20X water immersion lens (NA, 1.0).

### Quantification of cell and fibronectin density

Matlab was used to do the analysis. After immunostaining and imaging of large CAFs layers, the core of defects was automatically detected, and for each position, a circular mask of 400 pixels in diameter was taken. For each mask, the number of nuclei and the sum of pixel values inside the mask for fibronectin staining were calculated. The rest of the FOV was defined as no-defect areas. Both were then normalized to the area to obtain the cell and fibronectin density.

### Nematically aligned fibronectin network preparation and live imaging

To generate a nematically aligned fibronectin network, natural CAFs deposition of fibronectin was used. 6 x 10^4^ CAFs were seeded on 11 kPa PAA gels and cultured for 3 days in DMEM with 10% FBS, 1X ITS-A, 2X Anti-Anti, 12.5 µg/mL Metronidazole (B. Braun), and 4 µg/mL Ciprofloxacin (Panpharma) at 37°C 5% CO_2_. The entire CAF layer was imaged using brightfield on an Inverted Eclipse Ti-2 (Nikon) full motorized videomicroscope with a 10X objective (NA, 0.30). CAFs were then removed using 20 mM NH_4_OH and 0.1% Triton X-100 in PBS for 3 min at RT. PAA gels were washed several times with PBS and the fibronectin network was blocked for 15 min at RT, and incubated with the primary antibody for 30 min at RT. Following several washes in 0.05% Tween-20 in PBS, gels were incubated with the secondary antibody solution in blocking solution for 30 min at RT. After several washes, the entire gels were imaged using a videomicroscope (described earlier). PAA gels with the fibronectin network were kept in PBS with 12.5 µg/mL Metronidazole (B. Braun), and 4 µg/mL Ciprofloxacin (Panpharma) at 4°C until seeding with new CAFs. 60,000 CAFs were re-seeded onto the same gels coated with nematically aligned fibronectin for an additional 3 days of culture. Imaging of CAFs and fibronectin was repeated after these additional 3 days of culture, as performed before replating. For staining the fibronectin pattern after replating, a secondary antibody with a different Alexa Fluor was used to distinguish it from the initial fibronectin staining.

### Isotropic fibronectin network preparation and live imaging

To generate isotropic fibronectin layers, 2-3 x 10^5^ CAFs were seeded on 11 kPa collagen I-coated gels. After 1 day of culture, CAFs were removed using NH₄OH 20mM, 0.1% Triton-X-100 in PBS for 3 minutes at RT. Following several PBS washes, the remaining isotropic fibronectin matrix was stained (see Immunostaining). After imaging of the isotropic fibronectin pattern, 110,000 CAFs were seeded onto the same gels. Live imaging was performed starting 3 hours post-seeding and continued for 24 hours.

### Quantification of alignment correlation

Matlab and Fiji were used to do the analysis. The Fiji OrientationJ plugin ^47^ was used for each experiment to obtain the orientation maps θ of the CAFs and fibronectin layers. For each orientation map θ, the cosinus and sinus were calculated. The cosinus matrixes of CAFs and fibronectin were then multiplied together. Similarly, the sinus matrixes of CAFs and fibronectin were also multiplied together. Both results were summed together and the absolute value of it is taken. The mean of this absolute value matrix corresponds to the average alignment correlation between the CAFs and the fibronectin layer.

### Western Blot

CAFs were seeded on 30 kPa dishes (ExCellness - PrimeCoat, Biotech SA) coated with 5 µg/mL of monomeric collagen and treated with either control siRNA (siCtrl) or siRNA against fibronectin (siFN). After three days, cells were scratched and transferred to a falcon with PBS. Cells were washed twice with PBS by centrifugation for 2 min at 100 rcf. Cells were then re-suspended in 50 µL of Precellys and sonicated for 15s three times. Samples were boiled at 100°C for 10 min, then spun for 15 min at 15 900 rcf, and supernatants were diluted at one to five. Samples were then boiled at 95°C for 5 min and the same concentration of each sample was loaded into a 7,5% TGX gel and let migrate for about 1h. The proteins from the gel were transferred to a nitrocellulose membrane using Transbloc. The membrane was then incubated in 5% non-fat milk powder diluted in PBS on a shaker at RT for 1 h. The membrane was cut and then incubated at 4°C overnight with primary antibodies diluted in 1% milk/PBS solution on a shaker. The membrane was then washed three times for 5 min with PBS while shaking. The membrane was then incubated at RT for 1 h with the secondary antibodies conjugated with HRP. The membrane was then washed three times for 5 min with PBS while shaking. Cut membranes were put back together and the signal was revealed using an ECL substrate and visualized using a Chemidoc Touch Biorad.

### Drugs and siRNA

All drugs used in this study are listed in Table 3. Hoechst was added just before the acquisition: CAFs were washed with PBS and then incubated with Hoechst in PBS for 30 min at 37°C, 5% CO_2_. After incubation, CAFs were washed twice with PBS, cell medium was added, and acquisition was started. For siRNA treatment, CAFs were cultured for three days with siRNA on 30 kPa plates before being transferred on PAA gels. CAFs were then incubated for three more days with siRNA before launching image acquisition with siRNA present in the medium.

**Table 3.**
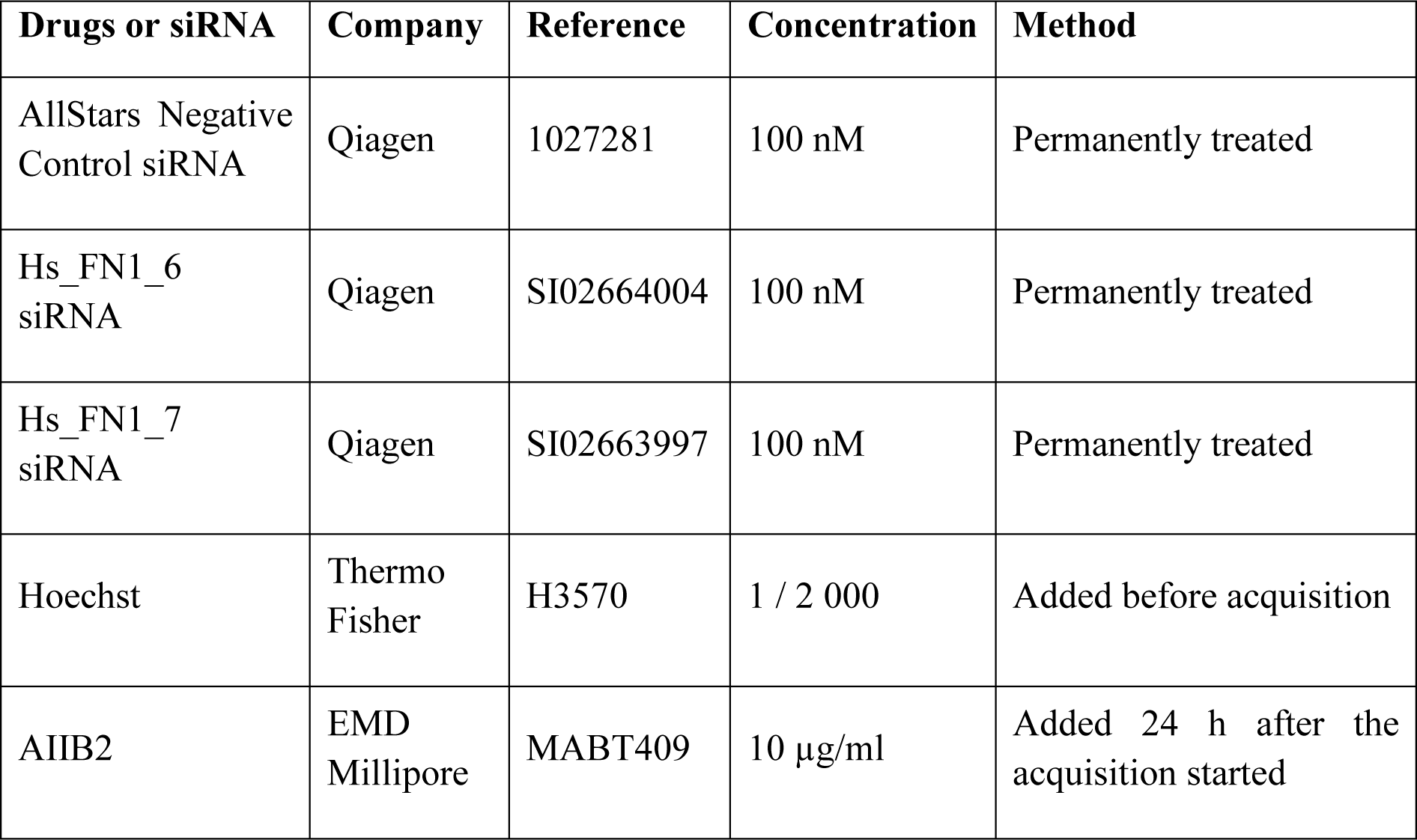
List of the drugs and siRNA used during this study.

### Defect trajectory analysis

The Fiji OrientationJ plugin was used for each experiment to obtain the orientation maps θ of the CAFs layers at each time point. For each experiment, positions were separated into different types of defects: comet and triangle defects. Defect cores were chosen to follow defect trajectory and found using the local nematic order parameter ***Q***. For each time point, the local nematic order parameter ***Q*** is calculated from the orientation maps θ using the following formula: ***Q***=√⟨𝑐𝑜𝑠2𝜃⟩^2^ + √⟨𝑠𝑖𝑛2𝜃⟩². The defect core was then found from the Q map as its minimum. Once the position of the defect core at each time point was obtained, translations of the trajectory coordinates were done to have the first time point of each core at (0,0) in the (x,y) axis. Velocity and persistence were calculated from each trajectory, using the @msdanalyser ^48^ of Matlab for persistence. Finally, all trajectories were plotted together for each type of topology, and boxplots for the velocities and persistence were obtained.

### Cell trajectory analysis

Cells’ trajectory was obtained by using Hoechst to stain nuclei or mixing a small proportion of LifeAct-GFP expressing CAFs with non-labeled CAFs and performing time-lapse imaging. For each position, Fiji was used to apply a threshold on images with the fluorescence of interest to generate binary images where each cell was well separated from the others. Thresholded images were then uploaded into Ilastik for each position to track cell trajectories. Each cell’s trajectory was saved into a file. Using Matlab, translations of cell trajectory coordinates were then done to have the first-time point of each cell at (0,0) in the (x,y) axis. Velocity and persistence were extracted from each trajectory, using the @msdanalyser ^48^ of Matlab for persistence. Finally, all cell trajectories were plotted together as well as boxplots for the velocities and persistence obtained.

### Mean square displacement analysis

All trajectories were analyzed using @msdanalyser ^48^ in Matlab.

### Quantification of the orientation and the fibronectin density for cell trajectories

CAFs were transfected with either control siRNA (siCtrl) or siRNA against fibronectin (siFN) and seeded onto preformed fibronectin patterns (see “Nematically aligned fibronectin network preparation”) at a density of 6 x 10^4^ CAFs per gel. 2 days post-seeding, nuclei were stained with Hoechst, and cells were imaged for 16 h with a 15 min interval (see “Time-lapse imaging of nematic ordering evolution over time”). In this case, multiple fields of view were acquired without stitching.

Images were processed using Fiji software. Briefly, the Hoechst signal was thresholded to obtain binary images of nuclei, and cell trajectories were computed using the Trackmate plugin^49^.

#### Orientation

In the Display option window, *Display spots* was unselected, but *Display tracks* was selected using the *Show entire tracks* option using a single color. The *capture overlay* action [select the first frame and hide the image] was then used to obtain a binary image of all trajectories present in the field of view. Using the *Analyze Particle* tool, trajectories were added to the ROI manager and enlarged by 10 pixels. These enlarged ROIs were then used to get a mask of the fibronectin layer present below the tracks; everything else in the field of view was filled with black. Finally, the orientation maps were obtained and calculated as described in “Quantification of alignment correlation”.

#### Fibronectin density

In the Display option window, *Display tracks* was unselected, but *Display spots* was selected. Using the TrackScheme, all initial spots were selected, and their corresponding position were exported using the *Export spots to IJ ROIs* action [export selection]. These ROIs were then used to measure the mean gray value on the fibronectin channel. All previous actions were repeated for the final spots.

### Quantification of the evolution of the fluid flow

Using Matlab, the velocity field of the whole CAF layer was calculated using the Particle Image Velocity (PIV) method. The PIV was used on the bright-field images of the whole CAF layer. The displacement in CAF position from one timepoint to the next one was measured on one-hundred-by-one-hundred-pixel squares along all images with overlapping between each square to precise the measure. From the displacement and time interval between each image, the velocities were calculated in each square, giving rise to the velocity fields at each time point. Finally, for each time point and condition, the means of the velocity fields were taken and plotted over time to show the evolution of the fluid flow.

