## Supplementary Information for "Aging and freezing of active nematic dynamics of cancer-associated fibroblasts by fibronectin matrix remodeling"

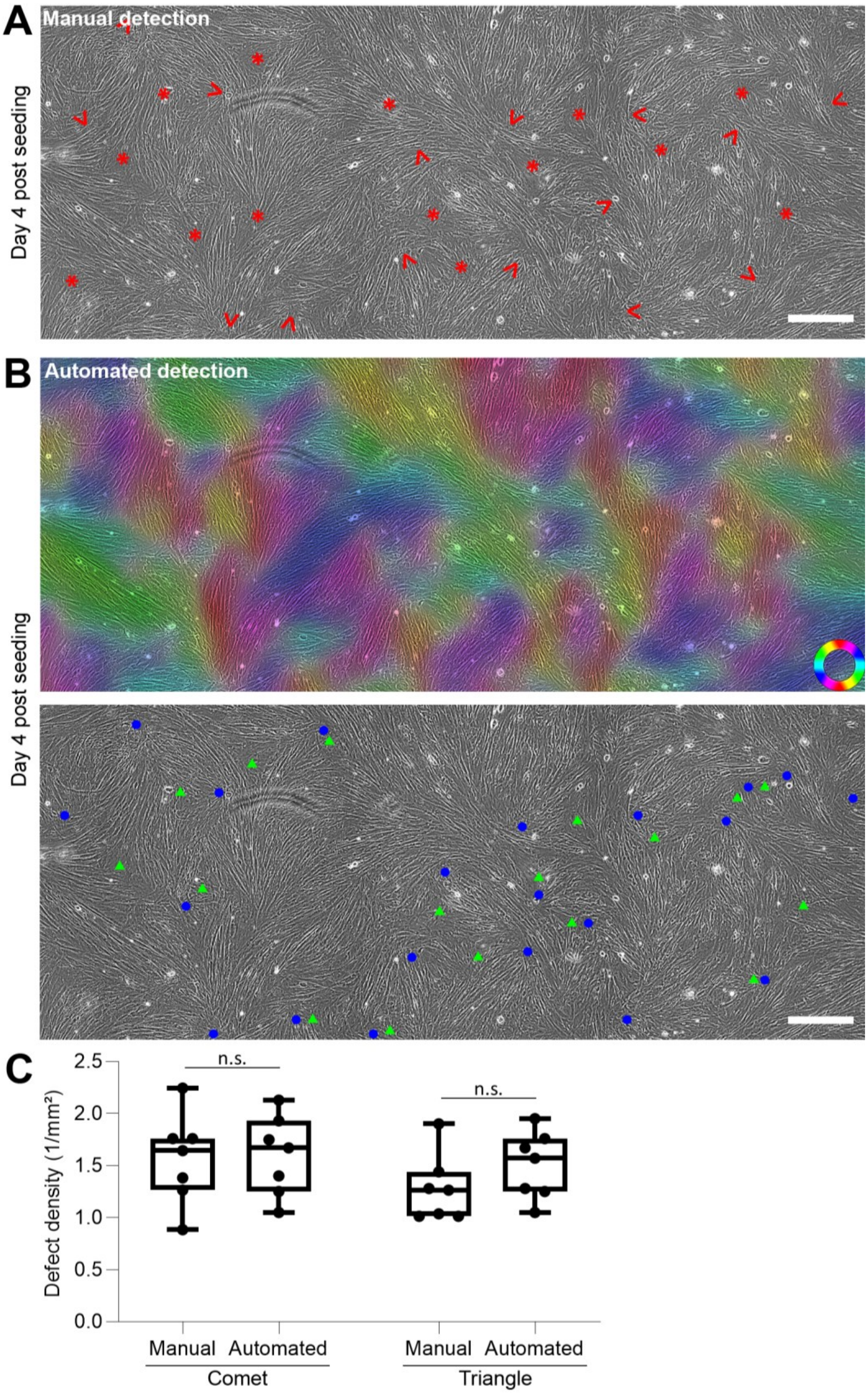


**Supplementary Figure 1. Comparison of manual and automated detection of defects in CAF layers**

A) Bright field of a CAFs layer cultured on an 11 kPa PAA gel for 4 days. Nematic order is perturbed by topological defects: manual detection of +1/2 or comet defects (arrowheads pointing towards comets' heads) and -1/2 or triangle defects (stars at the triangles' centers). Scale bar: 400 µm.

B) Top: orientation of CAFs based on the bright field image from panel A. Colored circle: orientation colormap.

Bottom: automated detection of topological defects: +1/2 or comet defects (blue dots) and -1/2 or triangle defects (green triangles). Scale bar: 400 µm.

C) Defect density in the nematically ordered CAF layer determined manually (panel A) or automatically (panel B). Comet and triangle defects from 7 fields of view. Each dot represents one field of view from 3 independent experiments. Boxplot: middle bar= median, edges bars= 25th and 75th percentiles, whiskers= extent of data. One-way ANOVA, p-value = 0.9981 and 0.6617, respectively.

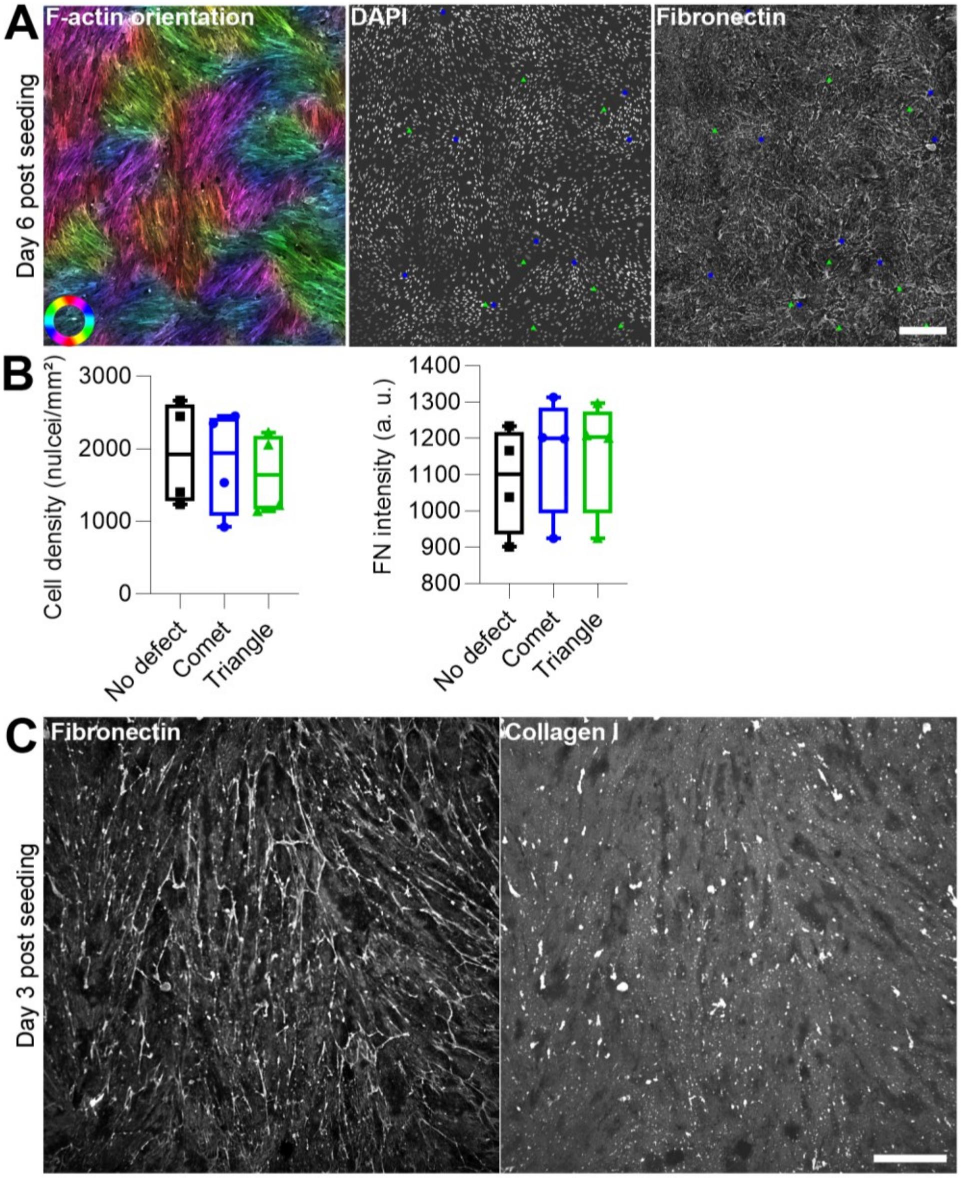


**Supplementary Figure 2. Cell and fibronectin density in the nematically ordered CAF layers**

A) Orientation of CAFs based on F-actin staining. CAFs nuclei (DAPI) and the fibronectin they deposited were imaged. Comet and triangle defects are represented as blue dots and green triangles, respectively. Colored circle: orientation colormap. Scale bar: 100 µm.

B) Quantification of local cell density and fibronectin density in the no-defect area, comet, and triangle defects. Each dot represents one FOV, data from one experiment. Boxplot: middle bar= median, edges bars= 25th and 75th percentiles, whiskers= extent of data.

C) Fibronectin and collagen I deposited by a CAF layer 3 days post seeding. Cells were removed before staining. Scale bar: 100 µm. Representative images of 2 independent experiments.

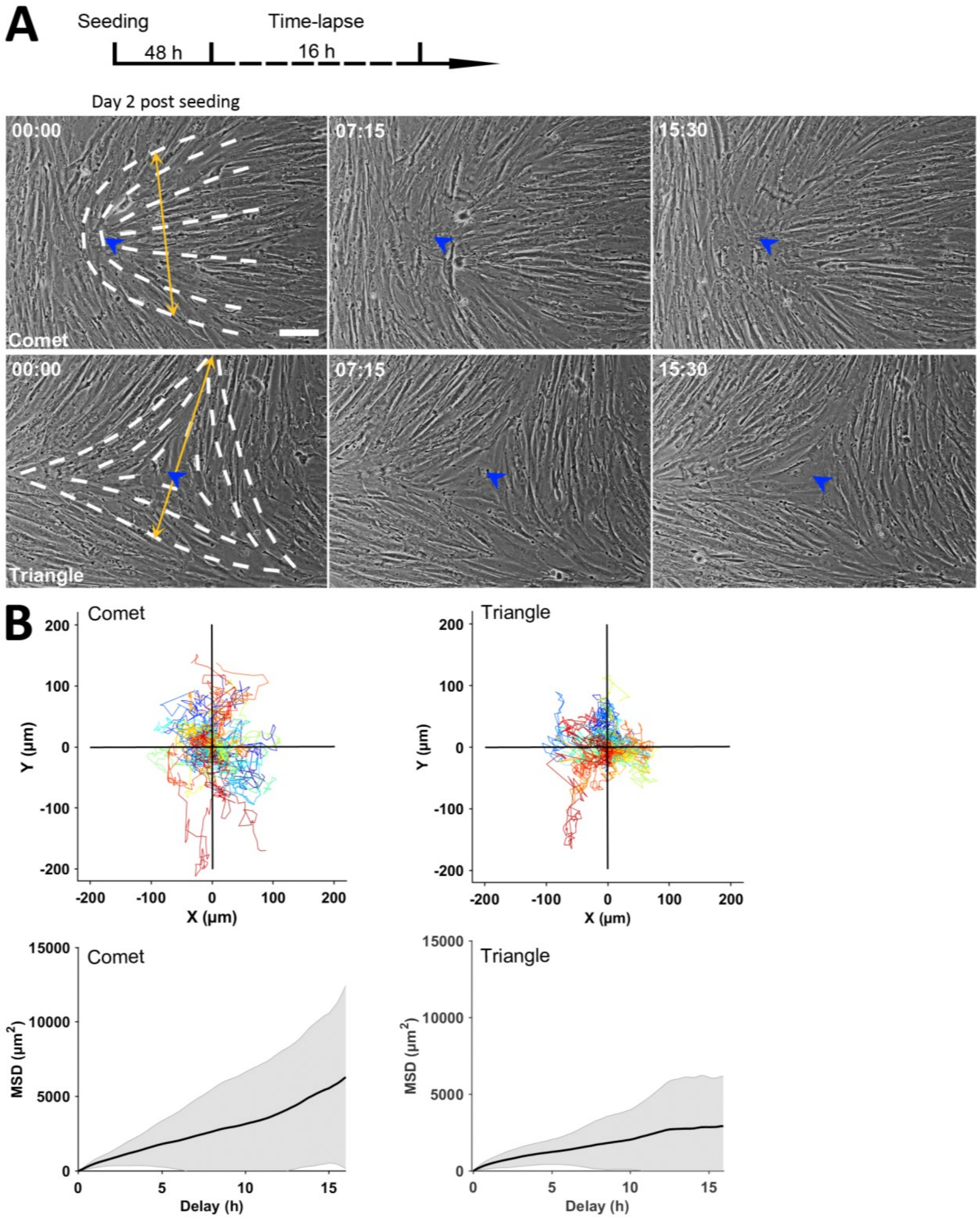


**Supplementary Figure 3. Manual tracking of defects in the CAF layer**

A) Time-lapse imaging of comet (top) and triangle (bottom) defect dynamics on a bright field 2 days after seeding. Time, hours:minutes. Scale bar: 100 µm. White dashed lines highlight the shapes of defects. Yellow double-headed arrows show the size of the defects. Blue arrowheads indicate the positions of defects' cores over time.

B) Trajectories of the defects' cores over the length of the time-lapse (top) and mean square displacement (MSD, bottom). For trajectories, horizontal and vertical black lines represent the defect size (400 µm). Comet (left): 35 and triangle (right): 31 defect trajectories from 3 independent experiments. The weighted average over all MSD curves is represented as a black line, and the weighted standard deviation over all MSD curves is represented as a gray area.

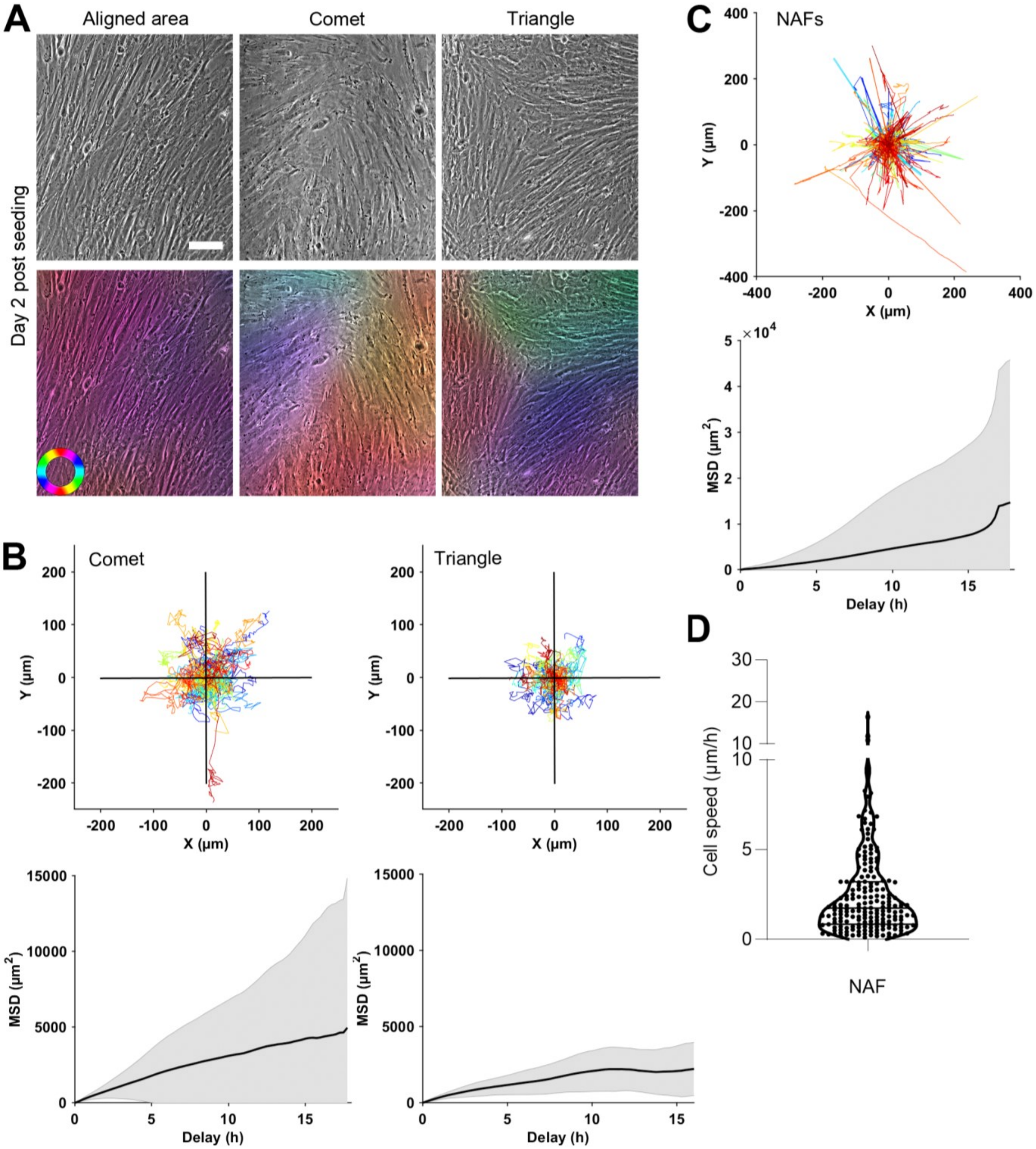


**Supplementary Figure 4. Defect and cell dynamics in NAF (normal associated fibroblast) layers**

A) Bright field of a NAF layer cultured on an 11kPa PAA gel for 2 days, showing aligned area (left) and topological defects: comet (middle) and triangle (right). Bottom images show the orientation colormaps. Colored circle: orientation colormap. Scale bar: 100 µm.

B) Trajectories of the defects' cores tracked manually over 16 h (top) and mean square displacement (bottom). Comet (left): 45 and triangle (right): 28 defect trajectories from 3 independent experiments. For trajectories, horizontal and vertical black lines represent the defect size (400 µm). The weighted average over all MSD curves is represented as a black line, and the weighted standard deviation over all MSD curves is represented as a gray area.

C) NAFs trajectories (top) and mean square displacement (bottom). 194 trajectories from 2 independent experiments. The weighted average over all MSD curves is represented as a black line, and the weighted standard deviation over all MSD curves is represented as a gray area.

D) Quantification of NAFs velocity. Each dot represents one NAF, 194 cells from 2 independent experiments.

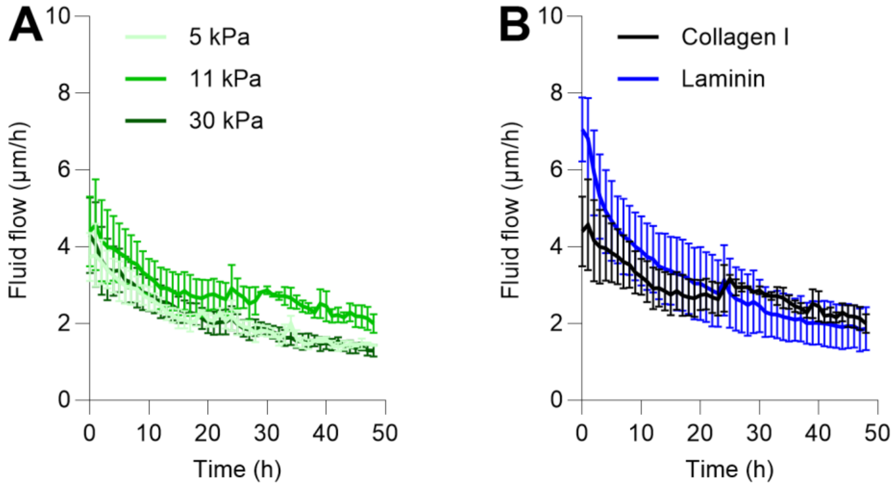


**Supplementary Figure 5. Freezing of cellular flows on different substrate stiffnesses and ECM coating**

A) Evolution of the mean of the velocity field of CAF layers plated on PAA gels of different stiffnesses over time. The stiffness of the PAA gel is represented in different shades of green (darker for stiffer). The lines represent the mean of 3 independent experiments, and the error bars represent the standard error of the mean.

B) Evolution of the mean of the velocity field of CAF layers plated on PAA gels coated with collagen I (black) or laminin (laminin-111; blue) over time. The lines represent the mean of at least 2 independent experiments, and the error bars represent the standard error of the mean.

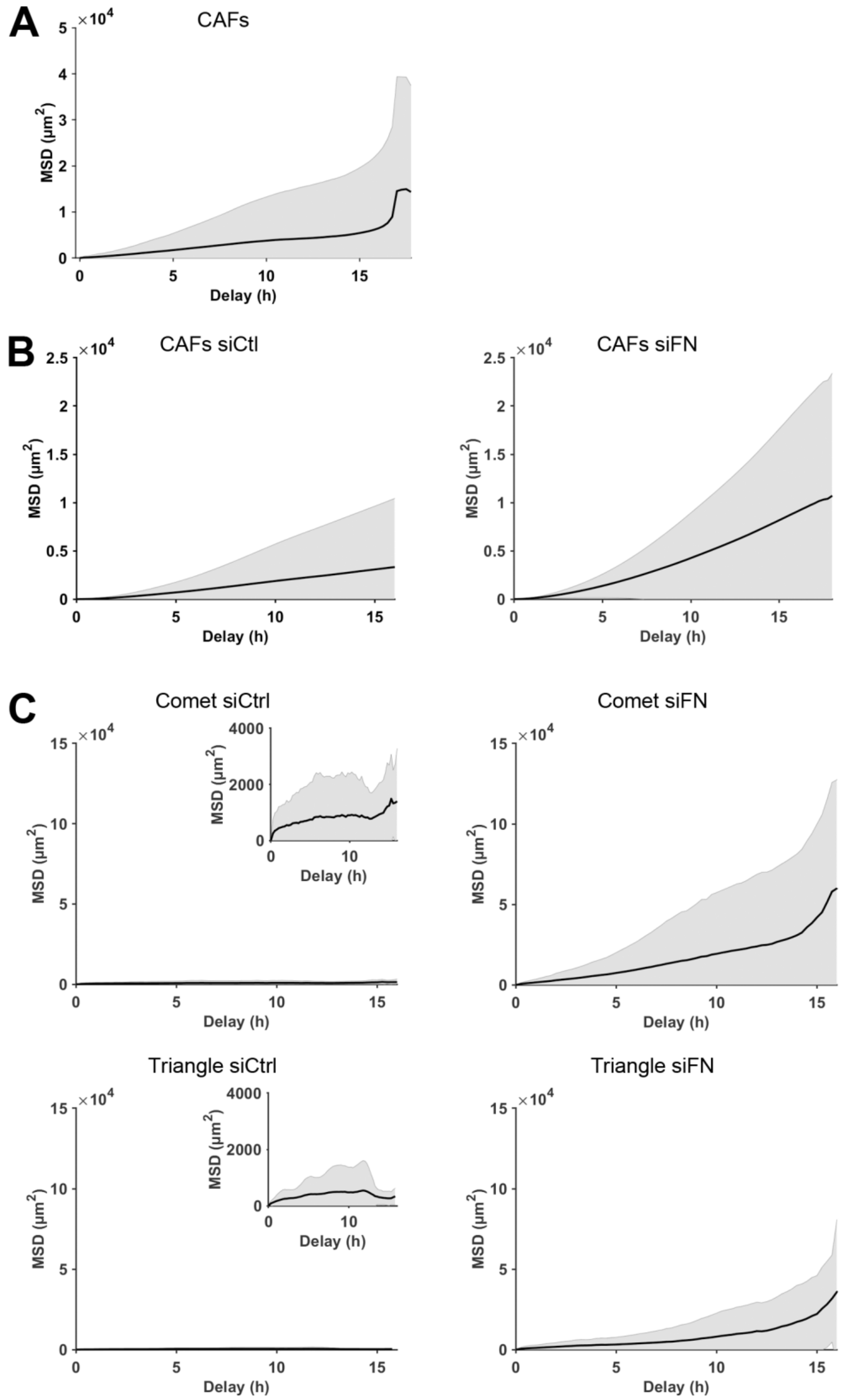


**Supplementary Figure 6. Mean square displacement for trajectories in Figures 2 and 3**

A) Mean square displacement (MSD) of CAFs, corresponding to the trajectories in Figure 2B.

B) Mean square displacement (MSD) of CAFs transfected with either control siRNA (siCtrl) or siRNA against fibronectin (siFN), corresponding to the trajectories presented in Figure 3C.

C) Mean square displacement (MSD) of defect cores in control (siCtrl) and fibronectin-depleted (siFN) CAFs layers, corresponding to the trajectories presented in Figure 3F. Insets show different scaling of the MSD axis for siCtrl defects.

The weighted average over all MSD curves is represented as a black line, and the weighted standard deviation over all MSD curves is represented as a gray area.

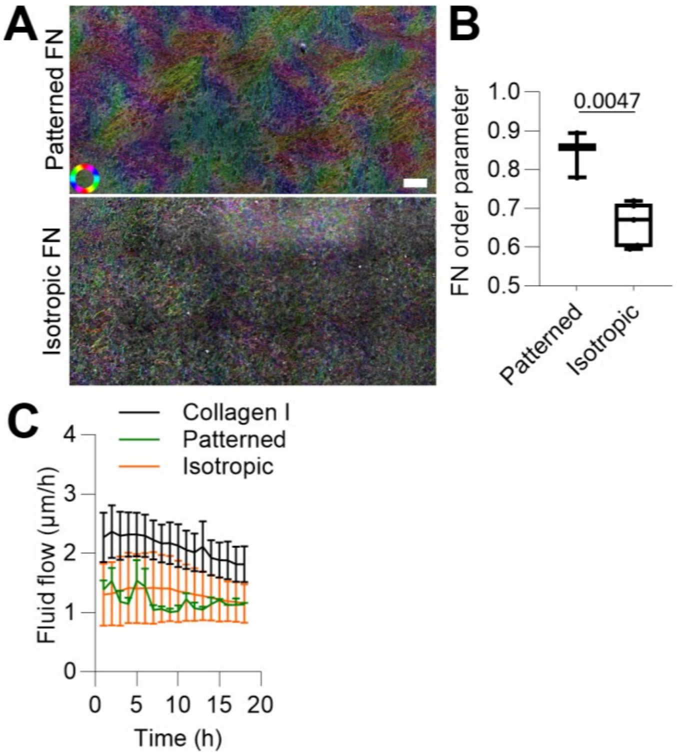


**Supplementary Figure 7. Effect of fibronectin alignment on CAF layer dynamics**

A) Orientation maps of fibronectin (FN) deposited by CAFs over 3 days (top, patterned FN) or over 1 day (bottom, isotropic FN). Colored circle: orientation colormap. Scale bar: 500 µm.

B) Quantification of the order parameter of fibronectin (FN) deposited by CAFs over 3 days (patterned) or over 1 day (isotropic). Data from at least 2 independent experiments; patterned: 3 gels; isotropic: 5 gels; two-tailed unpaired t-test. Boxplot: middle bar= median, edges bars= 25th and 75th percentiles, whiskers= extent of data.

C) Evolution of the mean of the velocity field of CAF layers plated on PAA gels coated with collagen I (black) or fibronectin deposited by CAFs over 3 days (patterned; green) or over 1 day (isotropic; orange) over time. The lines represent the mean 3 independent experiments, and the error bars represent the standard error of the mean.

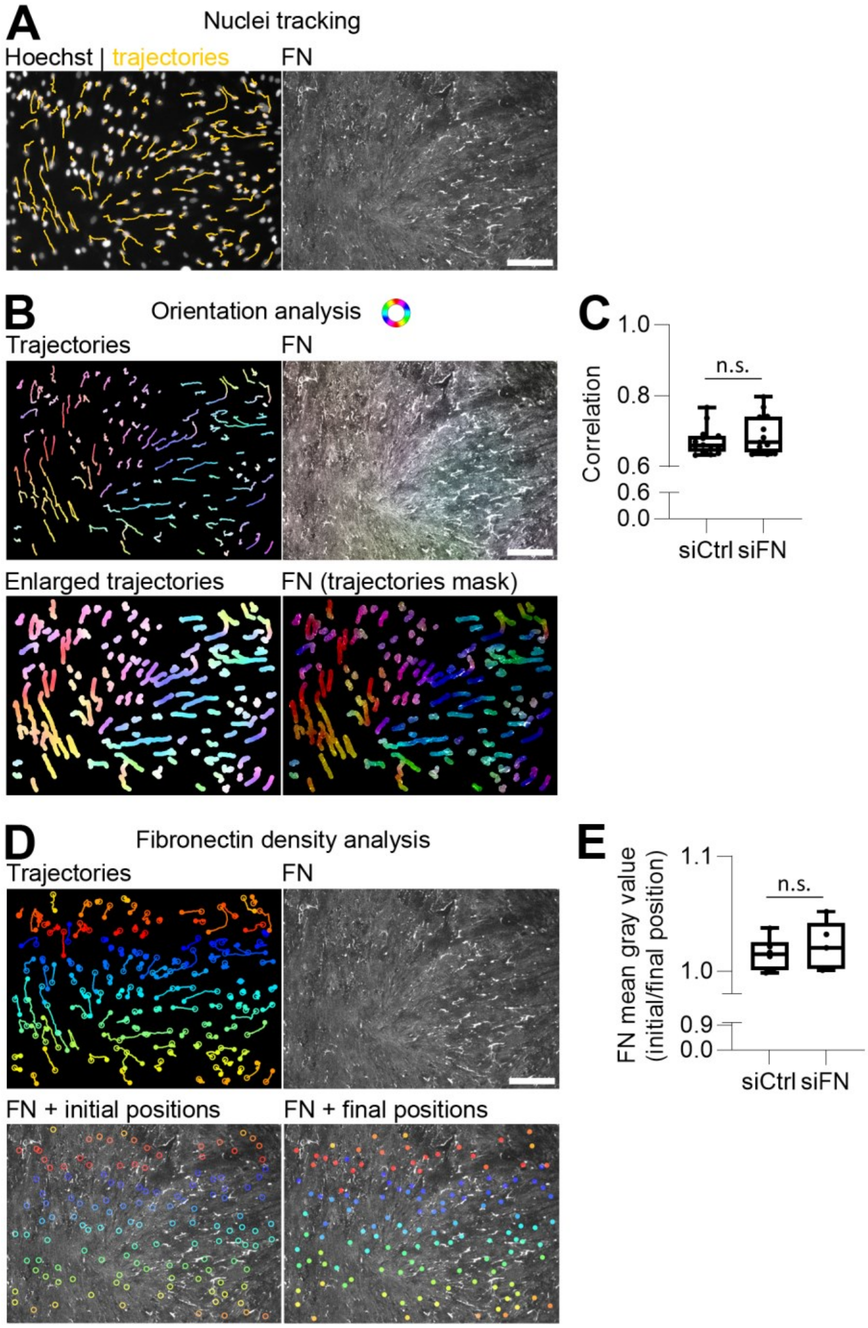


**Supplementary Figure 8. Orientation and fibronectin density analysis for trajectories in Figure 4F.**

A) Trajectories (yellow) of CAFs labeled with Hoechst (gray) 2 days after seeding on preformed fibronectin (FN) patterns. CAFs were tracked for 16h. Scale bar: 100 µm.

B) Orientation maps of trajectories and fibronectin (FN). Trajectories were enlarged by 10 pixels and used as a mask to compute the orientation of FN below them. Colored circle: orientation colormap. Scale bar: 100 µm.

C) Mean of the local spatial orientation correlation between the enlarged trajectories and the fibronectin layer below the enlarged trajectories. CAFs were transfected with either control siRNA (siCtrl) or siRNA against fibronectin (siFN). Each dot represents one field of view with about 100 trajectories; siCtrl: 15 fields of view; siFN: 13 fields of view from 3 independent experiments; two-tailed unpaired t-test: p-value = 0.2709. Boxplot: middle bar= median, edges bars= 25th and 75th percentiles, whiskers= extent of data.

D) Trajectories (one color per trajectory) of CAFs 2 days after seeding on preformed fibronectin (FN) patterns. For each trajectory, the initial and final position is represented as a circle and a dot, respectively.

E) Fibronectin (FN) mean gray value as a ratio of the initial and final position for a given trajectory. CAFs were transfected with either control siRNA (siCtrl) or siRNA against fibronectin (siFN). Each dot represents one field of view; siCtrl: 6 fields of view and 745 trajectories; siFN: 5 fields of view and 574 trajectories from 2 independent experiments; two-tailed unpaired t-test: p-value = 0.5385. Boxplot: middle bar= median, edges bars= 25th and 75th percentiles, whiskers= extent of data.

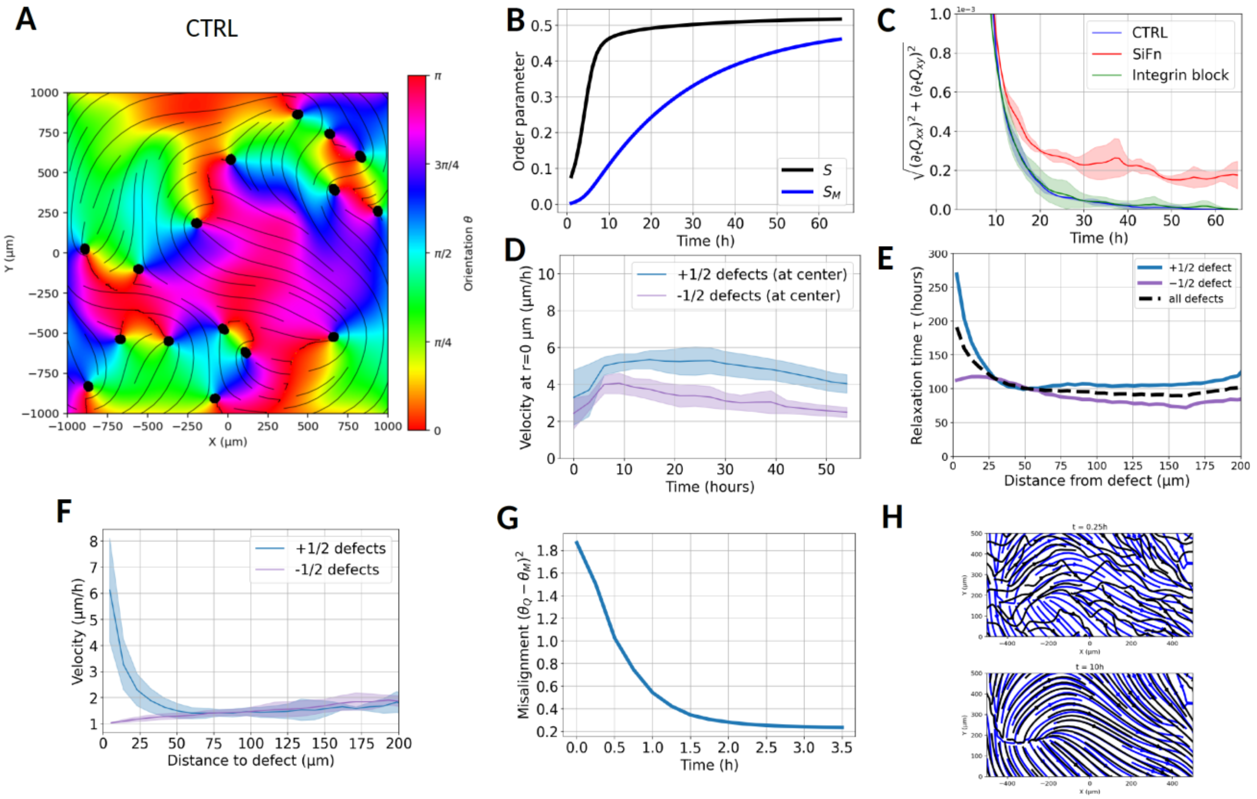


**Supplementary Figure 9. Extended analysis from the theoretical model**

Results B, C, D, E, and F are averaged over three simulations with three random initial conditions.

A) Orientational color map for a control condition after three days, showing CAF defect positions. Compare this to Figure 1C. The defect positions are indicated by black dots.

B) Aging of the nematic order parameter of the matrix and cell layer over time in the control case.

C) Freezing of the orientational pattern with time. The observable is the mean absolute value of the time derivative for the tensor order parameter Q.

D) Evolution of the fluid flow velocity averaged over the defect core for the +½ and −½ defects over time in the control case.

E) Relaxation time of the velocity as a function of distance from the +½ and −½ defects during aging of the control case.

F) Mean velocity profile of the fluid flow velocity versus distance from the defects after three days in the control case.

G) Relaxation of angle misalignment over time in prepatterned simulations.

H) Snapshots of nematic orientations for cells (black) and fibronectin (blue) at the beginning and after 10 hours of the prepatterned simulations.

### Supplementary model to
### "Aging and freezing of active nematic dynamics of cancer-associated fibroblasts by fibronectin matrix remodeling"

by Jacques et al.

#### I. MODEL DEFINITION

Cancer-associated fibroblasts (CAFs) are described as active apolar particles, which form a 2-dimensional layer of active nematic fluid; the corresponding nematic order parameter tensor is denoted by $\mathsf{Q} = S(\mathbf{nn} - \mathbf{I}/2)$, where $S$ is the amplitude of the order parameter and $\mathbf{n} \equiv (\cos\theta, \sin\theta)$ is the director field; it is a unit vector along the local direction of order parameterised by the angle field $\theta$ measured from an arbitrarily defined $\mathbf{x}$ axis. CAFs lay down apolar fibronectin fibres on their substrate, characterized by a nematic tensor $\mathsf{M} = S_M(\mathbf{mm} - \mathbf{I}/2)$, where $S_M$ is the strength of fibre field and $\mathbf{m} \equiv (\cos\theta_M, \sin\theta_M)$ is the director field corresponding to $\mathsf{M}$, parametrised by the angle field $\theta_M$. We construct below a minimal model of the coupled spatiotemporal dynamics for the CAFs and the fibre fields.

We model the dynamics of the active nematic (CAF) field $\mathsf{Q}$ as

$$\frac{D\mathsf{Q}}{Dt} = \frac{\mathsf{H}}{\gamma} - \lambda\mathsf{E} + \alpha_M\mathsf{M}. \tag{1}$$

Here, $D\mathsf{Q}/Dt = \partial_t\mathsf{Q} + (\mathbf{v}\cdot\nabla)\mathsf{Q} - (\mathsf{Q}\cdot\mathbf{\Omega} - \mathbf{\Omega}\cdot\mathsf{Q})$ is the corotational convective derivative where $\mathbf{\Omega} = [\nabla\mathbf{v} - (\nabla\mathbf{v})^T]/2$ is the vorticity; $\gamma$ is the rotational viscosity and $\mathsf{H} = -\delta F/\delta\mathsf{Q}$ is the molecular field that corresponds to the passive restoring force deriving from a free energy $F$ that penalizes deformations of the nematic field:

$$F = \int_\Omega d\mathbf{r}\left(\frac{K}{2}(\nabla\mathsf{Q})^2 + \frac{g}{4}(1 - 2(\mathsf{Q}:\mathsf{Q}))^2\right); \tag{2}$$

$\mathsf{E} = [\nabla\mathbf{v} + (\nabla\mathbf{v})^T - (\nabla\cdot\mathbf{v})\mathbf{I}]/2$ is the traceless strain-rate tensor and flow-alignment effects are parametrized by $\lambda$; last, the cell-matrix alignment interaction is introduced as an active aligning torque parametrized by $\alpha_M > 0$. This torque acts to align the CAF field $\mathsf{Q}$ parallel to the matrix $\mathsf{M}$.

The dynamics of the CAF layer is controlled by the friction with the substrate (with a friction coefficient $\xi$), its viscosity $\eta$, active stress $\zeta\mathsf{Q}$, and elastic stresses deriving from $F$ (since, in practice, the lengthscale $\sqrt{\eta/\xi}$ is expected to be small compared to all other lengthscales in our system—including $\sqrt{K/g}$—the viscosity will not feature in our theoretical discussions; it is included in our numerical simulations). The force balance equation is

$$\xi \mathbf{v} = -\nabla P - \zeta \nabla \cdot \mathbf{Q} + \eta \nabla^2 \mathbf{v} - \mathbf{H} : \nabla \mathbf{Q} + \lambda \nabla \cdot \mathbf{H} + \nabla \cdot (\mathbf{Q} \cdot \mathbf{H} - \mathbf{H} \cdot \mathbf{Q}) , \tag{3}$$

where $\mathbf{H} : \nabla \mathbf{Q}$ is $H_{kl} \nabla_i Q_{kl}$ and $P$ a pressure that enforces the constraint of incompressibility $\nabla \cdot \mathbf{v} = 0$. The friction $\xi$ may be isotropic, but later we will consider anisotropic friction $\xi(\mathbf{M})$, which will be a rank-2 tensor field that is a function of the matrix field $\mathbf{M}$. We have suppressed other active force densities at the same order in gradient ($\propto \mathbf{Q} \cdot \nabla \cdot \mathbf{Q}$) [1] since they do not affect our results qualitatively.

Finally, we need an equation for the dynamics of the matrix field $\mathbf{M}$. We consider a general case in which fibress are deposited in alignment with the CAFs, and can also degrade:

$$\partial_t \mathbf{M} = k \mathbf{Q} - k_d \mathbf{M}, \tag{4}$$

where $k$ and $k_d$ are respectively the deposition and degradation rates for the matrix field, which are uniform, and do not depend on cellular orientation.

Note that we also considered an alternative, phenomenological mechanism yielding a saturation of the field $\mathbf{M}$, which does not involve matrix degradation. This can be written:

$$\partial_t \mathbf{M} = k(S_0 - \mathrm{Tr}[\mathbf{M}^2]) \mathbf{Q}. \tag{5}$$

Finally, equations (1) to (3) above, together with either (4) or (5), fully define the coupled dynamics of the fields $\mathbf{Q}$ and $\mathbf{M}$. We discuss below striking features of the model.

#### II. SINGLE TOPOLOGICAL DEFECT DYNAMICS

In this section, we will discuss the matrix-induced freezing of active defect dynamics.

Our model is an extension of classical models of active nematic, which are known to display $\pm 1/2$ topological defects [2–5]. Indeed, our experiments display an abundance of $\pm 1/2$ defects in the CAF and matrix fields. In this section we use the dynamical description of Eq.(1)-(4) to examine how the topological defects in the $\mathbf{Q}$ field (i.e. the CAF layer) are affected by the coupling to the matrix $\mathbf{M}$ field. Since nematic $+1/2$ defects induce a polar director configuration, they spontaneously move in active systems [2–5]. We thus focus on the dynamics of $+1/2$ defects; we calculate their spontaneous velocity in our system using a method developed by Mazenko and Halperin [6] for equilibrium systems—and applied to active nematics in [7]—and show analytically how it depends on the coupling $\alpha_M$ (in the regime $\alpha_M$ small) to the matrix field.

A general calculation of the speed of an isolated $+1/2$ defect is complicated by the fact that the defect shape is modified by the active coupling to $\mathsf{M}$. However, if we assume that for small $\alpha_M$ the shape of the defect is not appreciably changed by this interaction, the only effect of (the defect in) $\mathsf{M}$ on the defect in $\mathsf{Q}$ is to change its speed. For this calculation, it is convenient to rescale lengths by the nematic coherence length, $\hat{\mathbf{r}} = \mathbf{r}/\sqrt{K/2g}$, and rescale time by the nematic relaxation time, $\tau = t/(\gamma/2g)$. This implies a scaling of the fluid velocity $\hat{\mathbf{v}} = \mathbf{v}/(\sqrt{2Kg}/\gamma)$. We further define complex coordinates $z = \hat{x} + i\hat{y}$ and complex order parameters $\psi_Q = Se^{2i\theta}$ and $\psi_M = S_M e^{2i\theta_M}$. The velocity of a defect at $z = z_0$ in the $\mathsf{Q}$ field is generically given by [6]:

$$v_Q = \left. \frac{-\partial_{\bar{z}}\bar{\psi}_Q\partial_\tau\psi_Q + \partial_{\bar{z}}\psi_Q\partial_\tau\bar{\psi}_Q}{\partial_z\psi_Q\partial_{\bar{z}}\bar{\psi}_Q - \partial_z\bar{\psi}_Q\partial_{\bar{z}}\psi_Q} \right|_{z=z_0}, \tag{6}$$

where the bar represents complex conjugation. Similarly, the velocity $v_M$ of a defect in the $\mathsf{M}$ field is given by the same equation with $\psi_Q$ replaced by $\psi_M$.

The rescaled dynamical equation for $\mathsf{Q}$ can then be written in terms of $\psi_Q$ as:

$$\partial_\tau\psi_Q + u\partial_z\psi_Q + \bar{u}\partial_{\bar{z}}\psi_Q = \lambda\partial_{\bar{z}}u + (\partial_z u - \partial_{\bar{z}}\bar{u})\psi_Q + 4\partial_z\partial_{\bar{z}}\psi_Q + (1 - |\psi_Q^2|)\psi_Q + \lambda_M\psi_M, \tag{7}$$

where $u = \hat{v}_{\hat{x}} + i\hat{v}_{\hat{y}}$, and $\lambda_M = \alpha_M\gamma/2g$. The dynamical equation for $\mathsf{M}$ in terms of $\psi_M$ is:

$$\partial_\tau\psi_M = \kappa(\psi_Q - (k_d/k)\psi_M), \tag{8}$$

where $\kappa = k\gamma/2g$.

We assume that the active and passive fields both have the form of an isolated $+1/2$ defect aligned in the same direction. We take the solution of the defect with a core at $Z_Q$ to be given by $\mathsf{H} = 0$ at all points in space, with $\theta$ having the appropriate winding ($\theta$ is not defined at the position of the defect). This is the shape that the defect has in the absence of all active terms (i.e., in the absence of $\alpha_M$ and $\zeta$). We also assume that the $\psi_M$ field has a defect of exactly the same form, but rescaled by the factor $k/k_d$ present in the deposition/degradation dynamics, at $z_M$ and calculate the velocity of the $\psi_Q$ defect due to this while assuming that defect shape remains unaffected for small $\alpha_M$ and $\zeta$ (this is a standard assumption usually made to determine speed of active defects). The defect solutions are:

$$\psi_Q = S(|z - z_Q|)\left(\frac{z - z_Q}{\bar{z} - \bar{z}_Q}\right)^{1/2}, \tag{9}$$

$$\psi_M = (k/k_d)S(|z - z_M|)\left(\frac{z - z_M}{\bar{z} - \bar{z}_M}\right)^{1/2}. \tag{10}$$

Since $S \sim r$ at short scales (where $r$ is the distance from the defect core), i.e., $S \sim |z - z_Q|$,

$$\psi_Q \propto z - z_Q, \tag{11}$$

$$\psi_M \propto (k/k_d)(z - z_M). \tag{12}$$

This yields the defect velocities for the active defect ($v_Q$) and the matrix defect ($v_M$):

$$v_Q = v_0 - \lambda_M (k/k_d)(z_Q - z_M), \tag{13}$$

$$v_M = \kappa (z_Q - z_M), \tag{14}$$

where $v_0$ is the flow velocity at the defect core of the $\mathbf{Q}$ field. In the small-deformation limit, we can find co-moving defect solutions $v_Q = v_M = v$:

$$v = \frac{v_0}{1 + \lambda_M k/(k_d \kappa)}, \tag{15}$$

with an inter-defect distance given by $z_Q - z_M = v/\kappa$. In dimensional units, this is given by:

$$v = \frac{v_0}{1 + \alpha_M / k_d}, \tag{16}$$

and the inter-defect distance is $|\mathbf{r}_Q - \mathbf{r}_M| = v/k$.

In the absence of matrix, the flow velocity at the defect core is, for fully compressible flow, given by the active flow:

$$v_0 = 2\alpha \partial_z \psi_Q = 2\alpha, \tag{17}$$

where $\alpha = -\zeta\gamma/K\xi$. For fully incompressible flow, this value is smaller: $v_0 = 3\alpha/2$ [7]. Our numerical results (Fig. 5B of main text) show that the prediction of Eq. (16) for the scaled velocity $v/v_0$ captures its dependence on $\alpha_M/k_d$. This shows that the deposition of a matrix by the active nematic field slows down a $+1/2$ defect's speed relative to a fast-relaxing matrix by a quantity that is controlled by the ratio of the time of defect remodeling, $1/k_d$, and of nematic relaxation to the matrix alignment, $1/\alpha_M$. In particular, the velocity of a $+1/2$ defect tends to zero as matrix remodeling (and in particular degradation) becomes infinitely slow, $k_d \to 0$.

#### III. FLOWS IN FROZEN ACTIVE NEMATIC TEXTURES

The above section shows that the velocity of a $+1/2$ defect vanishes in the limit $k_d \to 0$. To ensure that $\mathbf{M}$ also reaches a finite stationary state it is convenient to also assume $k \to 0$. In this limit, $\mathbf{M}$ can be considered as an external static field. We analyze static solutions of the $\mathbf{Q}$ field, and argue that active flows can be non-vanishing in this regime, in which both $\mathbf{M}$ and $\mathbf{Q}$ are frozen. Next, we argue that this flow can be eliminated by an anisotropic, $\mathbf{M}$-dependent friction.

##### A. Isotropic friction

To discuss this question theoretically, we start from the nematic director dynamics given by Eq. (1) and the force balance equation Eq. (3). In this limit, we have shown that the speed of a single defect vanishes i.e., $\mathsf{Q}$ reaches a stationary state; we assume that $\partial_t \mathsf{Q}$ also vanishes (i.e., $\mathsf{Q}$ reaches a steady-state) for an imposed $\mathsf{M}$ which has a more complex structure as well. We now look for a solution for which $\mathbf{v} = 0$ in addition in this case; i.e., $\partial_t \mathsf{Q} = 0, \mathbf{v} = 0$. From Eq. (1) this can only be the case if $\mathsf{H} = -\alpha_M \gamma \mathsf{M}$. Now inserting this into the force balance (3), we find that:

$$0 = -\nabla P - \zeta \nabla \cdot \mathsf{Q} + \alpha_M \gamma \mathsf{M} : \nabla \mathsf{Q} - \lambda \alpha_M \gamma \nabla \cdot \mathsf{M} - \alpha_M \gamma (\mathsf{Q} \cdot \mathsf{M} - \mathsf{M} \cdot \mathsf{Q}). \tag{18}$$

For this equation to be satisfied, $\nabla P$ *must* balance other force densities in (18); but because $\nabla P$ is a gradient field, it can only do so if the other force densities are also curl-free. This however does not hold generically, for arbitrary choices of $\mathsf{M}$. This shows that in general the velocity field does not vanish even when the $\mathsf{Q}$ field is frozen. Indeed, we see such situation in our simulations (see Fig. 5D and Supp. Fig. 9C).

##### B. Anisotropic friction

The observed nematic order in the matrix is generically expected to induce an anisotropic friction with the CAF layer. Anisotropic friction can be encoded by a mobility tensor $\boldsymbol{\mu}$ relating the velocity to the force applied to the substrate, $\mathbf{f}$ (the RHS of Eq. (3)) that differs from identity:

$$\mathbf{v} = \boldsymbol{\mu} \cdot \mathbf{f}. \tag{19}$$

The mobility tensor is constructed phenomenologically so that when there is no matrix ($S_M = 0$), the mobility is isotropic ($\boldsymbol{\mu} = \mu_0 \mathbf{I}$), while for fully-developed matrix ($S_M = 1$), the mobilities parallel and perpendicular to the matrix director $\mathbf{m}$ are $\mu_0$ and $\mu_\perp$ respectively:

$$\mathbf{m}^T \boldsymbol{\mu} \mathbf{m} = \mu_0 \tag{20}$$

$$\mathbf{m}_\perp^T \boldsymbol{\mu} \mathbf{m}_\perp = \mu_0 - S_M(\mu_0 - \mu_\perp), \tag{21}$$

where $\mathbf{m}_\perp$ is the vector normal to the matrix director field $\mathbf{m}$. The limiting case where the friction perpendicular to the matrix direction is infinite is the case where $\mu_\perp \to 0$, and so there can be no movement perpendicular to the direction of the fibres, $\mathbf{v} \cdot \mathbf{m} = |\mathbf{v}|$. This mobility tensor can be inverted to obtain the friction tensor.

In the case of a single defect, $\mathsf{M}$ does not form closed loops. Since $\mathbf{v}$ must be parallel to $\mathsf{M}$ at every point, the vanishing of the flow at infinity, together with the absence of sources and sinks in the continuity equation, that at steady state the flow is vanishing everywhere $\mathbf{v} = 0$.

Let us now consider a population of defects. While friction anisotropy in the model is parametrized by $\mu_\perp/\mu_0$, it is useful to introduce its space average value $\langle \mathbf{m}_\perp^T \boldsymbol{\mu} \mathbf{m}_\perp / (\mathbf{m}^T \boldsymbol{\mu} \mathbf{m}) \rangle$, where $\mathbf{m}_\perp$ is the normal vector to the matrix director field $\mathbf{m}$. For a developed matrix field $\mathsf{M}$ this is approximately related to the defect density $\rho$ and core area $A$ by:

$$\left\langle \frac{\mathbf{m}_\perp^T \boldsymbol{\mu} \mathbf{m}_\perp}{\mathbf{m}^T \boldsymbol{\mu} \mathbf{m}} \right\rangle = \frac{\mu_\perp}{\mu_0} (1 - \rho A) + \rho A. \tag{22}$$

Thus, even in the case of $\mu_\perp \to 0$, the space averaged anisotropy is non vanishing because defect cores remain disordered ($S_M \simeq 0$), and thus yield a locally isotropic friction. Flows are thus non vanishing close to defects cores, as is indeed observed in numerical simulations. This finally shows that anisotropic friction can significantly diminish hydrodynamic flows, which however can remain non-zero close to defect cores or along closed loops of streamlines of $\mathsf{Q}$, as is indeed observed in numerical simulations (see Fig. 5D and Supp. Fig. 9G).

#### IV. DETAILS OF THE NUMERICAL ANALYSIS

The numerical analysis of the model was carried out using the Finite Element Method software COMSOL Multiphysics [8]. We wrote the set of equations 1-4 described above as a weak form in the Weak Form PDE physics of the software with a linear element order, and the mesh size was chosen to be of the order of the defect core size ($\sqrt{K/g}$), which is the smallest characteristic size of the system. Anderson acceleration was added for stabilization and acceleration of the computation.

After declaring of the independant variables: `Hxx,Hxy,Qxx,Qxy,vx,vy,Mxx,Mxy,p`, the equations for the incompressible case were introduced in the Weak Form Physics in the following way:

```
test(Qxx)*Hxx[kg/h^2]-(test(Qxx)*Hxx0+K*test(Qxxx)*Qxxx+K*test(Qxxy)*Qxxy)
test(Qxy)*Hxy[kg/h^2]-(test(Qxy)*Hxy0+K*test(Qxyx)*Qxyx+K*test(Qxyy)*Qxyy)
test(Hxx)*Qxxt+(test(Hxx)*vx*Qxxx+test(Hxx)*vy*Qxxy)[µm/h]-test(Hxx)*Gammaxx-test(Hxx)*Qxy*(vxy-vyx)[µm/h]
test(Hxy)*Qxyt+(test(Hxy)*vx*Qxyx+test(Hxy)*vy*Qxyy)[µm/h]-test(Hxy)*Gammaxy+test(Hxy)*Qxx*(vxy-vyx)[µm/h]
(XIxx*vx+XIxy*vy)*test(vx)[µm/h]-test(vx)*(Hxx*Qxxx+Hxy*Qxyx)[kg/h^2]-test(vxy)*(Qxx*Hxy-Qxy*Hxx)[kg/h^2]
          +eta*(test(vxx)*vxx+test(vxy)*vxy)[µm/h]+zeta*test(vx)*(Qxxx+Qxyy)+test(vx)*px[kg/h^2]
(XIyx*vx+XIyy*vy)*test(vy)[µm/h]-test(vy)*(Hxx*Qxxy+Hxy*Qxyy)[kg/h^2]+test(vyx)*(Qxx*Hxy-Qxy*Hxx)[kg/h^2]
          +eta*(test(vyx)*vyx+test(vyy)*vyy)[µm/h]+zeta*test(vy)*(Qxyx-Qxxy)+test(vy)*py[kg/h^2]
test(Mxx)*(Mxxt-GammaMxx)
test(Mxy)*(Mxyt-GammaMxy)
test(p)*(vxx+vyy)[µm]
```

while the independent variable `p` disappears in the compressible case. Integration by parts has been used the reduce the order of the derivatives in the equations. The polynomial parts of the molecular field write `Hxx0=2*g*Qxx*(-1+2*2*(Qxx^2+Qxy^2))`,`Hxy0=2*g*Qxy*(-1+2*2*(Qxx^2+Qxy^2))`. The CAFs nematic source terms are `Gammaxx=-muxx[kg/h^2]/gamma-alphaM/gamma*Mxx` and `Gammaxy=-muxy[kg/h^2]/gamma-alphaM/gamma*Mxy` where the flow alignment terms are neglected. Anisotropic friction is introduced as `XIxx=xi*(1-anis*(Sm/2+Mxx))/(1-anis*Sm)`,`XIxy=XIyx=-xi*(Mxy*anis) /(1-anis*Sm)`, and `XIyy=xi*(1-anis*(Sm/2+Myy))/(1-anis*Sm)`, where `Sm=sqrt(Mxx^2+Mxy^2)*2`.

The parameters of the numerical studies are `g,K,gamma,alphaM,xi,zeta,eta` are shown in Table.SI (we show the mobility which is the inverse of the friction). The anisotropy parameter `anis` varies between 0 (isotropic friction) and 1 (fully anisotropic). The settings of the numerical studies are as follows:

**Fig 5B**: We study the co-evolution of a cell layer $+1/2$ defect that is initially superimposed on a matrix $+1/2$ defect. We have free boundary conditions. The system is assumed to be compressible. The equations modelled in these figures are Eqs. 1-2-3 while the dynamical equation for the matrix is Eq. 4. The aim of this simulation was to compare the numerically obtained velocity with the analytical results and to show its dependence on the interactions between the matrix and the CAF layer. For this reason, no viscosity was introduced, as the analytical solution for the velocity is based on a momentum equation with only friction.

**Fig 5C**: We study the time evolution of an incompressible multidefect system containing both matrix and CAFs, with no cell growth. The initial conditions for the nematic orientation of the CAFs are random, while the initial conditions for the matrix are $S_M = 0$. The friction is anisotropic according to Eq. 19-21. The equations modeled in these figures are Eqs. 1-2-3 whereas the matrix follows Eq. 5 and does not degrade. The orientation of the cells is constrained to be parallel to the boundaries of the system. We focus in this figure on the impact of friction anisotropy on the fluid flow.

**Fig 5D-E**: We study the time evolution of an incompressible multidefect system containing both matrix and CAFs, with no cell growth, for CTRL, SiFn and integrin knocked out systems. The initial conditions for the nematic orientation of the CAFs are random, while the initial conditions for the matrix are $S_M = 0$. The friction is anisotropic in the CTRL case, and the equations modeled in these figures are Eqs. 1-2-3, whereas the matrix follows Eq. 5 and does not degrade. The orientation of the cells is constrained to be parallel to the boundaries of the system.

| Figure | b | d-e-f-g | Ref |
|---|---|---|---|
| $g$ $(kg.h^{-2})$ | $10^5$ | $10^4$ | $10^5\, kg.\mu m^{-1}.h^{-2}$ [9] |
| $K$ $(kg.\mu m^2.h^{-2})$ | $4\times 10^7$ | $10^7$ | $10^{5,6}\, kg.\mu m.h^{-2}$ [9] |
| $\alpha_M$ $(h^{-1})$ | $0.13 \to 0.4$ | 0.27 | current study |
| $\gamma$ $(kg.h^{-1})$ | $1.5\times 10^5$ | $1.2\times 10^4$ | $10^{4,5}\, kg.\mu m^{-1}.h^{-1}$ [9] |
| $k$ $(h^{-1})$ | $0 \to 100$ | $6.7\times 10^{-2}$ | current study |
| $\zeta$ $(kg.h^{-2})$ | $-2\times 10^5$ | $-2.1\times 10^6$ | $\pm 10^{3,4}\, kg.\mu m^{-1}.h^2$ [9, 10] |
| $\mu_{\parallel}(S_M=1)$ $(kg^{-1}.\mu m^2.h)$ | $2.5\times 10^{-3}$ | $5.5\times 10^{-4}$ | $10^{-1,-2}$[10] |
| $\mu_{\perp}(S_M=1)$ $(kg^{-1}.\mu m^2.h)$ | $2.5\times 10^{-3}$ | $0 \to 5.5\times 10^{-4}$ | $10^{-1,-2}$ [10] |
| $\eta$ $(kg.h^{-1})$ | 0 | $1.1\times 10^4$ | $10^{2-4}\, kg.\mu m^{-1}.h^{-1}$ [11–14] |
| $\ell_c = \sqrt{K/g}$ (core size, $\mu m$) | 20 | 31.6 | $\sim 2$ [9] |
| $\sqrt{\eta/\xi}$ (screening length, $\mu m$) | 0 | 2.5 | 10 [15, 16] |
| $\gamma/(g+\alpha_M\gamma)$ (rotational timescale, $h$) | $\sim$ | 0.9 | $0.1-1$ [9] |
| $\zeta/(\ell_c(1/\mu+\eta/\ell_c^2))$ (defect velocity, $\mu m.h^{-1}$ ) | $\sim 25$ | 6.0 | $1-10$ [17, 18] |

Table S I. Parameters used in the numerical studies. The parameters are written in the first column, and the values used for each numerical study are written in the next 3 columns. The last column gives references for the values we have introduced. Our study is in 2 dimensions, so the units are different from those used in the literature, which describe 3D systems. Note that for simplicity we have taken the flow alignment parameter to be zero $\lambda = 0$ in all numerical studies.

#### V. POST-PROCESSING OF THE NUMERICAL STUDY

**Fig 5B**: We localised the surface minimum value for $S_M$ and $S$ using the *non local coupling* in COMSOL Multiphysics. We then considered these coordinates as the centres of the matrix and the cell layer defects.

**Fig 5C**: The root mean square of the flow was extracted directly from COMSOL Multiphysics post-processing.

**Fig 5D-E**: Defect velocity was measured on day 6 of the experiment. Defect tracking was performed using ImageJ-win64 (Plugin TrackMate) [19, 20]. The movies acquired by COMSOL Multiphysics were transformed into binary images. Defects were then tracked with the Tracking/-TrackMate plugin.
